# MHCII reduction is insufficient to protect mice from alpha-synuclein-induced degeneration and the Parkinson’s HLA locus exhibits epigenetic regulation

**DOI:** 10.1101/2024.08.31.610581

**Authors:** Elizabeth M Kline, Janna E Jernigan, Christopher D Scharer, Jeffrey Maurer, Sakeenah Hicks SL, Mary K Herrick M, Rebecca L Wallings, Sean D Kelly, Jianjun Chang, Kelly B Menees, Nikolaus R McFarland, Jeremy M Boss, Malú Gámez Tansey, Valerie Joers

**Affiliations:** Department of Molecular and Cell Biology, University of Connecticut, Storrs, CT USA; Department of Neuroscience, University of Florida College of Medicine, Gainesville, FL USA; Center for Translational Research in Neurodegenerative Disease, University of Florida College of Medicine, Gainesville, FL USA; Department of Microbiology and Immunology, Emory University School of Medicine, Atlanta, GA USA; Department of Cell Biology, Emory University School of Medicine, Atlanta, GA, USA; Department of Neurology, University of Florida College of Medicine, Gainesville, FL USA; Norman Fixel Institute for Neurological Diseases, University of Florida Health, Gainesville, FL USA; McKnight Brain Institute, University of Florida Health, Gainesville, FL USA

**Keywords:** Antigen presentation, MHCII, microglia, Parkinson’s disease, T cells, alpha-synuclein, HLA

## Abstract

Major histocompatibility complex class II (MHCII) molecules are antigen presentation proteins and increased in post-mortem Parkinson’s disease (PD) brain. Attempts to decrease MHCII expression have led to neuroprotection in PD mouse models. Our group reported that a SNP at *rs3129882* in the MHCII gene Human leukocyte Antigen (HLA) DRA is associated with increased MHCII transcripts and surface protein and increased risk for late-onset idiopathic PD. We therefore hypothesized that decreased MHCII may mitigate dopaminergic degeneration. During an ongoing α-synuclein lesion, mice with MHCII reduction in systemic and brain innate immune cells (LysMCre+I-Ab^fl/fl^ or CRE+) displayed brain T cell repertoire shifts and greater preservation of the dopaminergic phenotype in nigrostriatal terminals. Next, we investigated a human cohort to characterize the immunophenotype of subjects with and without the high-risk *GG* genotype at the *rs3129882* SNP. We confirmed that the high-risk *GG* genotype is associated with peripheral changes in MHCII inducibility, frequency of CD4+ T cells, and differentially accessible chromatin regions within the MHCII locus. Although our mouse studies indicate that myeloid MHCII reduction coinciding with an intact adaptive immune system is insufficient to fully protect dopamine neurons from α-synuclein-induced degeneration, our data are consistent with the overwhelming evidence implicating antigen presentation in PD pathophysiology.

## Introduction

Evidence from multiple studies suggests an association between changes in the immune system and the hallmark neuropathologies of Parkinson’s disease (PD), specifically age-related dopaminergic degeneration and α-synuclein aggregation. The brain’s resident immune cells, microglia, are activated in rodent models of transgene- and viral-vector-mediated α-synuclein overexpression^1–5^, as well as in the post-mortem^6–9^ and living brains of individuals with PD^10,11^. Cytokines such as IL-1β, TNF, and IL-6 are secreted by activated microglia and promote an inflammatory environment in the brain, as seen in PD substantia nigra *pars compacta* (SN, locus of neurodegeneration), and striatum^12,13^. Cerebrospinal fluid and serum levels of inflammatory cytokines correlate with disease duration and motor symptom severity^14,15^, and can sensitively and specifically differentiate healthy controls from individuals with PD^16^. Adaptive immune system alterations have also been identified in PD, including extensive infiltration of CD4+ and CD8+ T cells into the central nervous system of individuals and rodent models^17,18^. Autoantibodies against dopaminergic (DA) neuron proteins in PD patient serum and cerebrospinal fluid have also been observed^19^, as have T cells capable of recognizing synuclein-derived antigens^20^. Despite what we have learned about the state of the immune system in PD, the mechanism by which immune system function influences risk for PD remains unclear.

One key immune system function of interest in PD is the process of antigen presentation, which is the nexus between the innate and adaptive immune systems. The way in which antigen presentation regulates the status of the immune system specifically in PD is not yet fully understood. In general, antigen presentation relies on major histocompatibility complex proteins (MHC) proteins which display antigen peptides to T cells, prompting T cell activation and differentiation. Differentiated T cells promote pathogen clearance and the formation of immunological memory. Microglia and other specialized antigen presenting cells display peptides derived from engulfed pathogens and intracellular debris using MHC class II (MHCII), recognized by CD4+ T cells. In *in vitro* and *in vivo* studies, microglia have been shown to have the capacity to phagocytose certain species of α-synuclein as well as debris released by degenerating DA neurons^21–24^. More recent work, however, highlights border associated macrophages (BAMs), a developmentally distinct population of brain-resident antigen presenting and phagocytic cells, as the key to PD-related neurodegeneration^25^. Genetic variations in the MHCII locus are associated with synucleinopathies and PD risk^26,27^. Coupled with the observation that MHCII expression in PD brain correlates with deposition of α-synuclein^28^ and is increased relative to control brains^6^, the current evidence suggests the possibility that immune cells phagocytose and present α-synuclein-derived peptides or antigens from α-synuclein-burdened, degenerating neurons to CD4+ T cells in PD. While the precise location of such antigen presentation (within the brain parenchyma, in the perivascular space, or more distant from the central nervous system) is a subject of debate, the fact that antigen presentation is an important process in PD is accepted.

Consistent with post-mortem observations and studies with animal models of PD in which MHCII is upregulated in the brain, a single nucleotide polymorphism (SNP) at *rs3129882* within the gene for MHCII, which in humans is called *Human Leukocyte Antigen* (*HLA*), was reported to be associated with a 1.7-fold increase in risk for idiopathic late-onset PD^29^. The high-risk genotype (*GG*) at *rs3129882* is over-represented (46%) in individuals with idiopathic PD compared to its frequency in an age-matched healthy control population (40%). Functionally, our group demonstrated that *GG* homozygosity at this SNP is associated with greater baseline mRNA and surface MHCII protein expression in monocytes and B cells^30^. We interpreted these findings to mean that the high-risk genotype might also be associated with increased antigen presentation throughout the lifespan, thereby increasing risk for PD as chronic antigenic load increases as a function of age^31^. It may be the case that genotype differences at *rs3129882* may also be associated with differences in T cell activation and differentiation. While a recent study reports that there are no differences between PD and controls in T cell activation in response to common antigens^32^, there are differences in T cell response to PD-specific antigens, like α-synuclein^20^. We hypothesize that *rs319882* genotype, given its impact on MHCII expression, may influence T cell function.

Because immune activation, including upregulation of MHCII, is associated with neurodegeneration, there has been interest in targeting MHCII for PD treatment, with the goal of blocking antigen presentation to prevent aberrant T cell activation and subsequent neurodegeneration. Previous work using a MHCII global knockout reported that MHCII and CD4+ T cells are necessary for α-synuclein-induced dopaminergic degeneration^33–35^. While these studies were informative, they examined the immune response in a context in which CD4+ T cells were not able to develop normally, complicating interpretation of the results. Indeed, some expression of MHCII is necessary to activate T cells when needed to fight an infection, for example, levels above the homeostatic range are linked to autoimmune diseases and accompanied by abnormal ratios of T cell subsets^36^. Therefore, our approach was to use a mouse model where MHCII is conditionally reduced from targeted myeloid lineages and determine the extent to which T cell frequencies and dopamine (DA) neuron integrity are altered. Towards this effort, we used a model of PD pathology in which a recombinant adeno-associated virus (rAAV) drives human α-synuclein expression in mouse SN. To understand the contributions of innate immune cell MHCII-mediated antigen presentation to α-synuclein-induced inflammation and degeneration, mice with lysozyme M (LysM)Cre-based reduction of MHCII (I-Ab^fl/fl^) were used in these experiments, as LysM is mainly expressed on myeloid cells (Figure 1). Furthermore, we characterized the relationship between high- and low-risk genotype at *rs3129882* and immunophenotype, specifically T cell subsets, in individuals with PD and control subjects.

**Figure 1.**
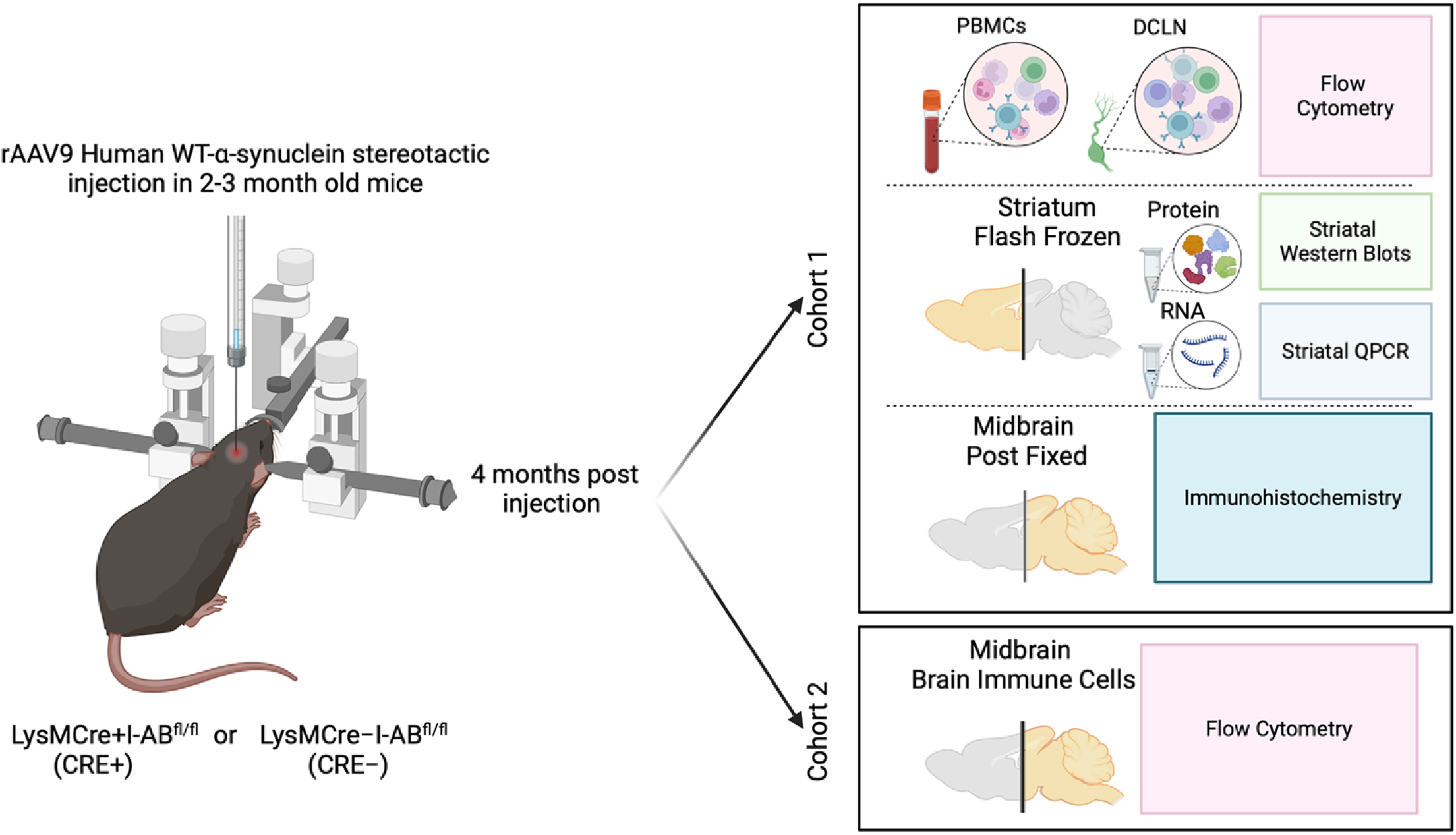
Experimental design and outcomes. Experiments were split between 2 cohorts of LysMCre+I-Ab^fl/fl^ (CRE+) or LysMCre-I-Ab^fl/fl^ (CRE-) mice. Both cohorts were given stereotaxic unilateral injections of rAAV2/9-human WT α-synuclein (rAAV9 hWT-ASYN) into the substantia nigra and euthanized 4 months later. Peripheral blood mononuclear cells (PBMCs) and deep cervical lymph nodes (DCLN) were collected for flow cytometry, striatum was microdissected and evaluated with Western blots and qPCR for immune markers, and caudal brain was post-fixed and evaluated using immunohistochemistry to verify AAV targeting. Cohort 2 had brains collected for flow cytometry analysis. Figure created with BioRender.com.

## Results

### Confirmation of myeloid cell subtype effects of targeted reduction of MHCII

The extent to which Cre expression under the LysM promoter affected brain MHCII protein expression was determined in LysMCre+I-Abfl/fl (CRE+) and LysMCre-I-Abfl/fl (CRE-) mice 4 months post-rAAV2/9 human WT α-synuclein (rAAV9 hWT-ASYN) nigral injection. Cohort 1 was assessed via immunohistochemistry and cohort 2 using flow cytometry. Quantification of MHCII-immunoreactive cells in the SN reveal significant reduction of MHCII-immunoreactivity in CRE+ mice relative to CRE- (Figure 2A-C). Additionally, brain immune cells evaluated by flow cytometry demonstrated a trend in reduced MHCII+ microglia and significant decreases in the MHCII mean fluorescent intensity (MFI) in microglia from CRE+ mice (Figure 2D-E). Furthermore, significant reductions were identified in MHCII frequency and MFI in monocytes (Ly6G-CD11c-) and neutrophils (LY6G+CD11c-) in CRE+ mice, while no differences were found in dendritic cells (Ly6G-CD11c+) when comparing genotypes (Figure 2F-K). In a separate cohort, naïve animals were evaluated for MHCII expression on immune cells from the spleen and peritoneal cavity. Like brain immune cells, these peripheral immune cell populations demonstrated robust reduction of MHCII on myeloid populations in CRE+ mice (Supplementary Figure S1A-F).

**Figure 2.**
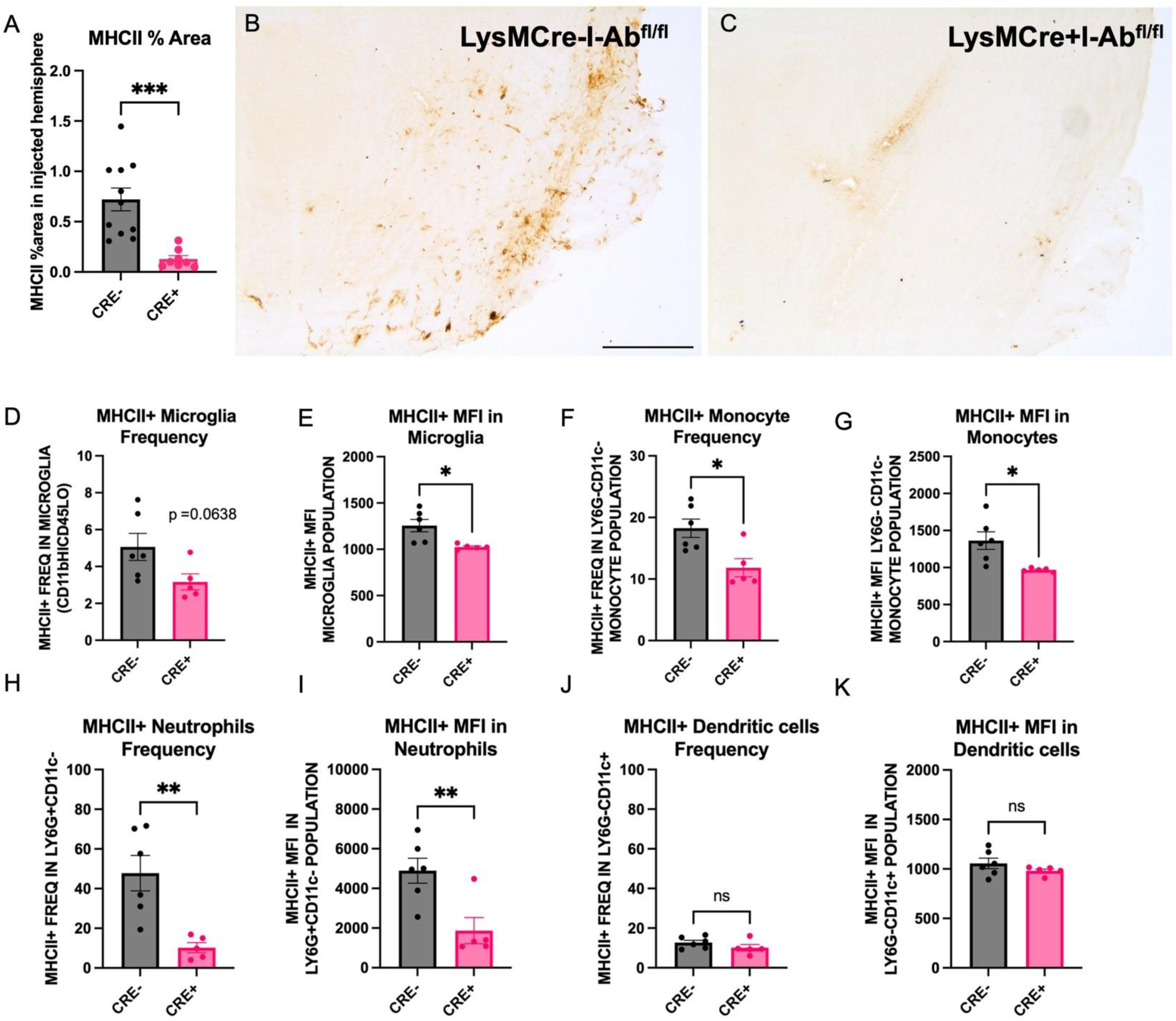
Successful reduction of brain MHCII in myeloid cells of CRE+ mice. (A) Quantification of MHCII-immunoreactivity in the midbrain of LysMCre-I-Ab^fl/fl^ (CRE-) and LysMCre+I-Ab^fl/fl^ (CRE+) mice demonstrate significant reduction in CRE+ animals. (B,C) Representative microphotographs of MHCII immunostaining in the midbrain. (D-K) Brain immune cells harvested by Percoll gradient and stained for flow cytometry analysis show reduced frequency of MHCII+ microglia, monocytes, and neutrophils, but not dendritic cells monocytes in CRE+ animals compared to CRE-. Mean fluorescent intensity (MFI) of MHCII+ cells was also reduced on microglia, monocytes, and neutrophils but not dendritic cells in CRE+ animals compared to CRE-. Each dot represents an individual animal and data is plotted as mean ± standard error of mean. *p<0.05,**p<0.01,***p<0.001. Unpaired two-tailed t-test (Welch’s correction was used for those measures with significantly different variances). Scale bar, 500μm.

### Reduction of myeloid MHCII in CRE+ mice has a global effect on CD4 T-cell subsets

To determine the effect of peripheral myeloid MHCII reduction on T cell subset frequency, peripheral blood mononuclear cells (PBMCs) were isolated from CRE- and CRE+ four months after stereotaxic injection with rAAV9 hWT-ASYN to the SN. We observed a significant increase in frequency of naïve CD4+ T cell (CD44-CD62L+) and decreased CD4+ frequencies of central memory (CD44+CD62L+), effector memory (chronically activated, CD44+CD62L-) and Tregs (Tbet-FoxP3+) in CRE+ mice (Figure 3A,E), with no changes in total CD4+ cells (Figure 3C). These results are consistent with the decreases in MHCII expression found in CRE+ mice, given that CD4+ T cell activation and differentiation is dependent on antigen presentation via MHCII. These significant effects were not found in PBMC CD8 populations (Figure 3B, D), although there was a trend to increase both frequency of total CD8 (p=0.059) and naïve CD8 T cells (p=0.0873). When evaluating lymphocyte populations in the brain (Figure 4A-D), similar effects of CD4 subsets were found in the brains of CRE+ mice, such that the frequency of CD4+ naïve cells (CD44-CD62L+) was increased and frequency of CD4+ central memory cells (CD44+CD62L+) was decreased (Figure 4E). In contrast to the PBMCs, CD8 lymphocytes were altered in the brain of CRE+ mice, such that the frequency of CD8+ cells were significantly increased and a trend to decrease CD8+ central memory cells (CD44+CD62L+; p=0.0524; Figure 4F). The deep cervical lymph nodes (DCLN) are the central nervous system-draining lymph nodes. In immune cells isolated from these small structures, we report a decreased frequency of CD4+ effector memory (CD44+CD62L-) and CD4+ Tregs (FoxP3+Tbet-; Figure 4G, H). The CD4+ T cell population shifts present throughout central and peripheral immune fractions in CRE+ suggest peripheral myeloid reduction of MHCII has a global effect on CD4+ T cell frequencies.

**Figure 3.**
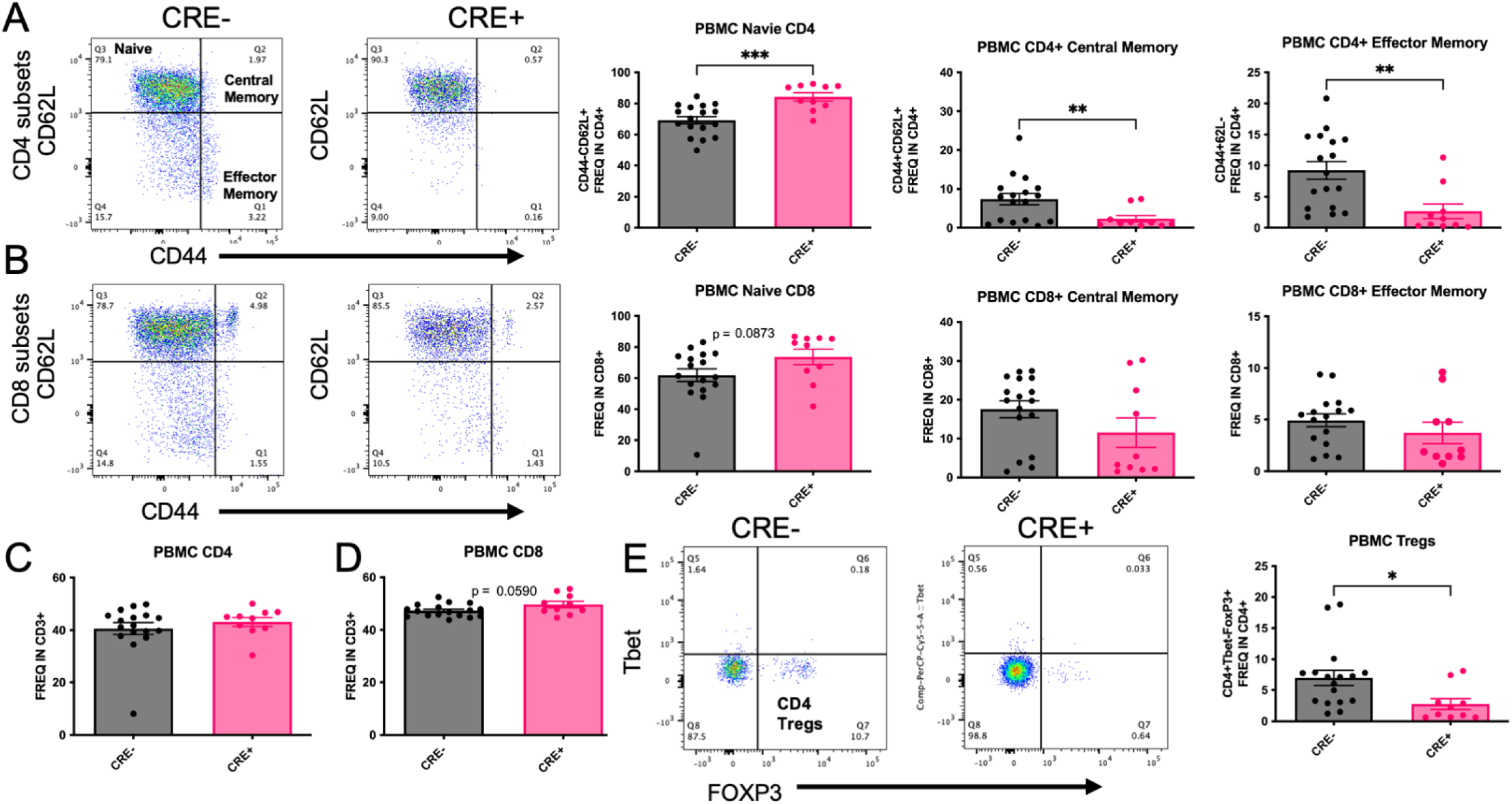
Reduction of myeloid MHCII in CRE+ mice has a global effect on CD4 T-cell subsets. PBMCs were isolated from LysMCre-I-Ab^fl/fl^ (CRE-) and LysMCre+I-Ab^fl/fl^ (CRE+) mice and evaluated for T cell subsets using flow cytometry. (A) Representative dot blots and quantifications of CD4+ T cell subsets show that naïve CD4+ T cells (CD44-CD62L+) were increased in CRE+ animals compared to CRE-, while CD4+ central memory (CD44+CD62L+) and CD4+ Effector memory (CD44+CD62L-) were decreased in CRE+. (B) Representative dot blots and quantifications of CD8+ T cell subsets show there was only a trend to increase (p=0.0873) in CD8+ naïve T cells, with no difference in CD8+ central memory and Effector memory cells. (C,D) Yet, frequencies of CD4 and CD8 T cells within the CD3+ population were not different between genotypes. (E) Representative dot blots and quantification display a decrease in Tregs (Tbet-FoxP3+) in CRE+ animals. Data are plotted as mean ± standard error of the mean (SEM). P values are indicated for unpaired two-tailed t-test (Welch’s correction was used for those measures with significantly different variances): *p<0.05, **p<0.01, ***p<0.001.

**Figure 4.**
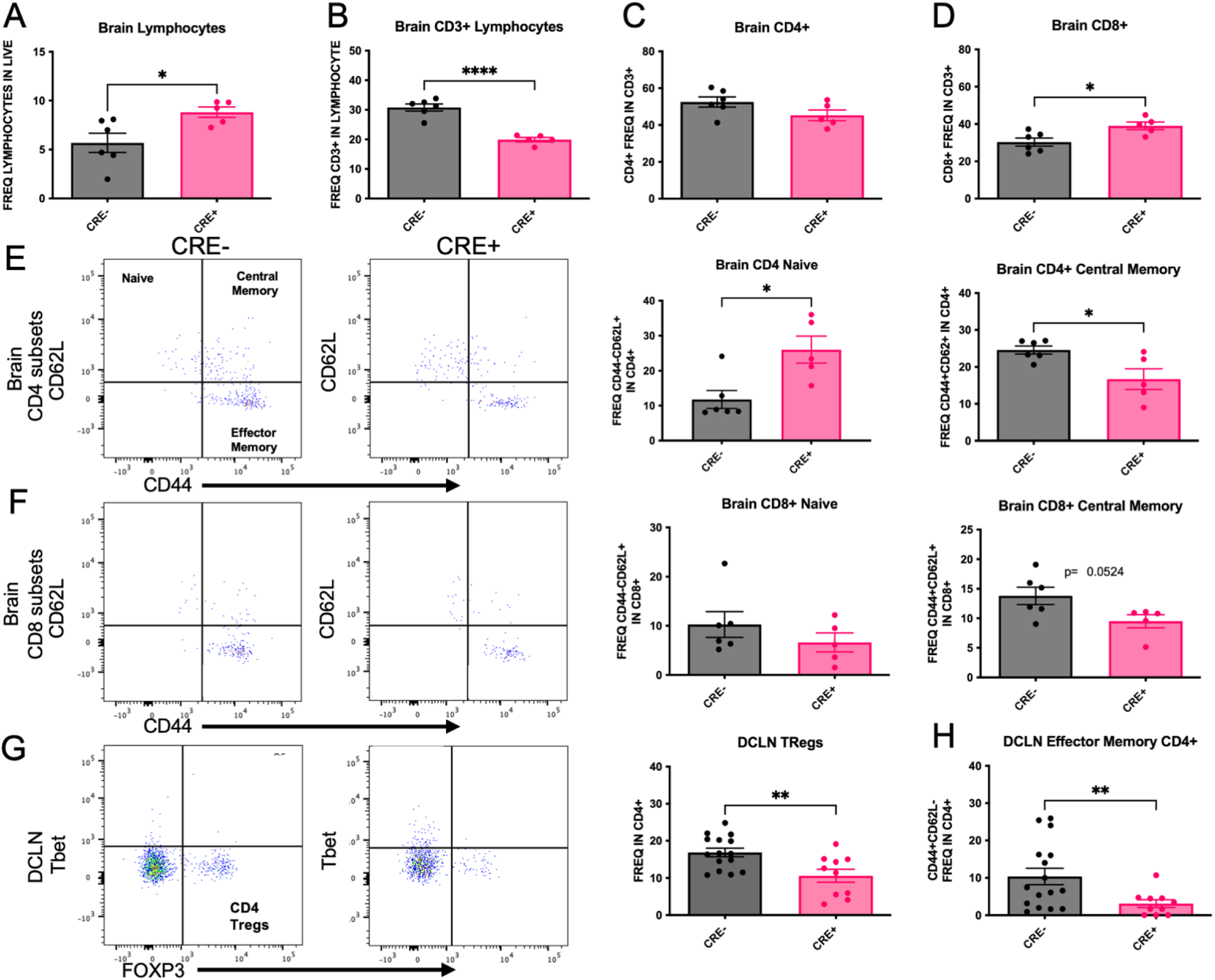
Brain and DCLN CD4+ T cell subsets are affected by LysMCre-mediated reduction of MHCII. Brain immune cells were isolated from LysMCre-I-Ab^fl/fl^ (CRE-) and LysMCre+I-Ab^fl/fl^ (CRE+) mice and evaluated by flow cytometry. (A,B) Total brain lymphocyte frequencies were increased in CRE+ animals compared to CRE-, while CRE+ animals exhibited decreased frequencies of CD3+ immune cells compared to CRE-. (C,D) CD4+ frequencies were the same in CRE+ and CRE-animals, and CRE+ displayed a statistically significant increase in CD8+ frequency compared to CRE-. Representative dot blots and quantification of (E) CD4+ and (F) CD8+ cell subsets in the brain find an increase in CD4+ naive (CD44-CD62L+) cells in CRE+ with no differences in CD8+ naïve. Brain CD4+ central memory (CD44+CD62L+) were decreased in CRE+ animals with a trend to decrease (p=0.0524) in CD8+ central memory brain frequencies. Immune cells from the left and right deep cervical lymph nodes (DCLN) were isolated from CRE- and CRE+ mice. (G) Representative dot blots and quantification of DCLN CD4+ Tregs and Th17 subsets show decreased in Tregs (FoxP3+) and (H) effector memory (CD44+CD62L-) in CRE+ animals. Data are plotted as mean ± standard error of the mean (SEM). P values are indicated for two-tailed t-test: *p<0.05, **p<0.01, ****p<0.0001.

### Striatal dopaminergic terminals from CRE+ mice are detectably protected 4 months after rAAV9 hWT-ASYN injection

The loss of TH expression in the striatum following nigral rAAV9 hWT-ASYN in mice has been reported elsewhere^37^. Here, we confirmed that in our hands, at a titer of 2.1×10^12^, rAAV9 hWT-ASYN virus compromised DA neuron phenotype in the striatum, as indicated by a significant decrease in DAT and TH protein expression in both normal expressing MHCII mice and MHCII-deficient mice (Figure 5A,B). Post hoc analysis controlling for multiple comparisons revealed that, in CRE+ animals, the striatum ipsilateral to virus injection had statistically significantly more DAT protein than in CRE- (Figure 5A). Furthermore, the post hoc analysis revealed CRE-striatal TH content was also significantly lower ipsilateral to the human α-synuclein compared to the uninjected striatum, while CRE+ mice were protected from this effect (Figure 5B). Furthermore, phosphorylated TH was decreased by viral vector-mediated α-synuclein overexpression in both genotypes (Figure 5C). These data and the representative western images (Figure 5D) suggest reduction of myeloid MHCII is associated with maintenance of DA neuron phenotype (DAT and TH expression, capacity to transport and synthesize dopamine) despite ongoing α-synuclein burden. Interestingly, there was a statistically significant effect of genotype on the extent of α-synuclein burden. We observed increased α-synuclein in SN (by immunofluorescence) and striatum (by western) on the side ipsilateral to viral vector injection in both genotypes, indicative of anterograde transport of human WT α-synuclein from the SN. Evaluation of striatal lysates determined that the increase in α-synuclein protein was greater in CRE-mice compared to CRE+ mice (Supplementary Figure S2).

**Figure 5.**
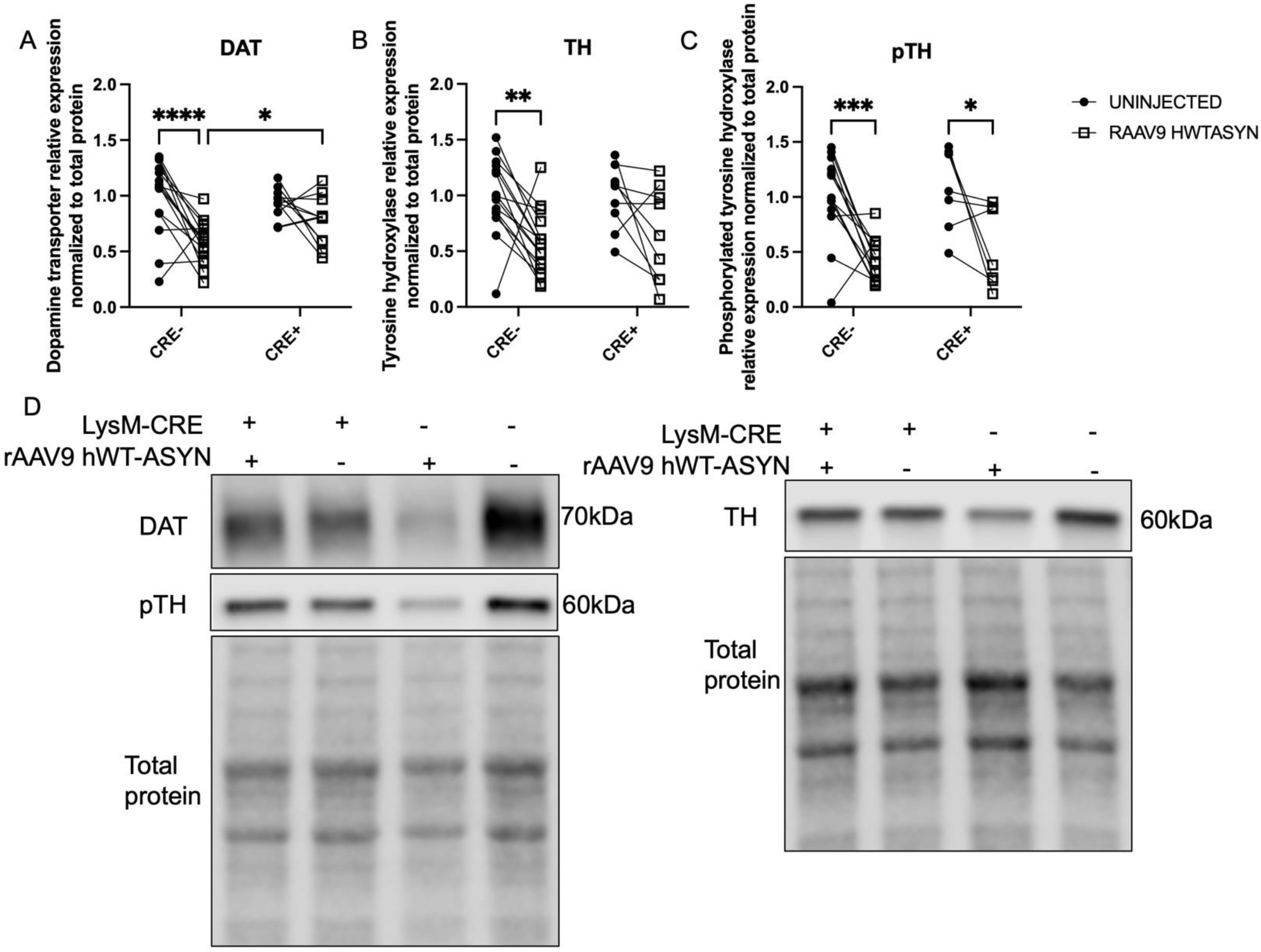
Striatal dopaminergic terminals from CRE+ mice are detectably protected 4 months after rAAV9 hWT-ASYN injection. Soluble striatal protein lysates from the uninjected hemisphere and rAAV2/9 human WT α-synuclein (RAAV9 hWT-ASYN) injected hemisphere of LysMCre-I-Ab^fl/fl^ (CRE-) and LysMCre+I-Ab^fl/fl^ (CRE+) mice were analyzed for protein levels of (A) dopamine transporter or DAT, (B) tyrosine hydroxylase or TH, and (C) phosphorylated TH (pTH) normalized to total protein and relative expression based on CRE-average to compare across blots. Data are plotted to show the paired striatal samples for each animal and evaluated with a two-way repeated measures analysis with an uncorrected Fisher’s LSD multiple comparisons. (D) Representative immunoblots of DAT (70kDa), pTH (60kDa) and TH (60kDa) staining and their respective total protein used for loading normalization. Gels are cropped from originals that can be found as supplemental data. *p<0.05, **p<0.01, ***p<0.001, ****p<0.0001.

### Striatal inflammatory responses of mice with targeted reduction in MHCII to alpha-synuclein is significantly dampened

The transcription of several immune markers was investigated to characterize the inflammatory status of the striatum, where nigral DA neurons project. We observed an effect of human α-synuclein gene expression such that multiple comparisons analysis found a significant increase between uninjected compared to injected striatum of CRE-mice (Figure 6A), indicating α-synuclein mRNA from the rAAV2/9 virus was present in the striatum. The striatum ipsilateral to the rAAV9 hWT-ASYN injection presented with increased CD68 (lysosomal marker) and IBA1 (myeloid marker) gene expression compared to the uninjected striatum in CRE-mice, but difference between hemispheres was not significant in CRE+ mice (Figure 6B-C). These results suggest mice with sufficient levels of MHCII have increased phagocytosis and myeloid cell burden (IBA1) in the brain in the presence of human WT α-synuclein, while those with reduced myeloid MHCII do not display the expected innate immune response. There were no statistically significant effects of either the viral injection or LysMCre genotype on CD4 (Figure 6D), TNF, CCL2, MHCII, MHCI, IFN-γ, and IL-1b mRNA level (data not shown).

**Figure 6.**
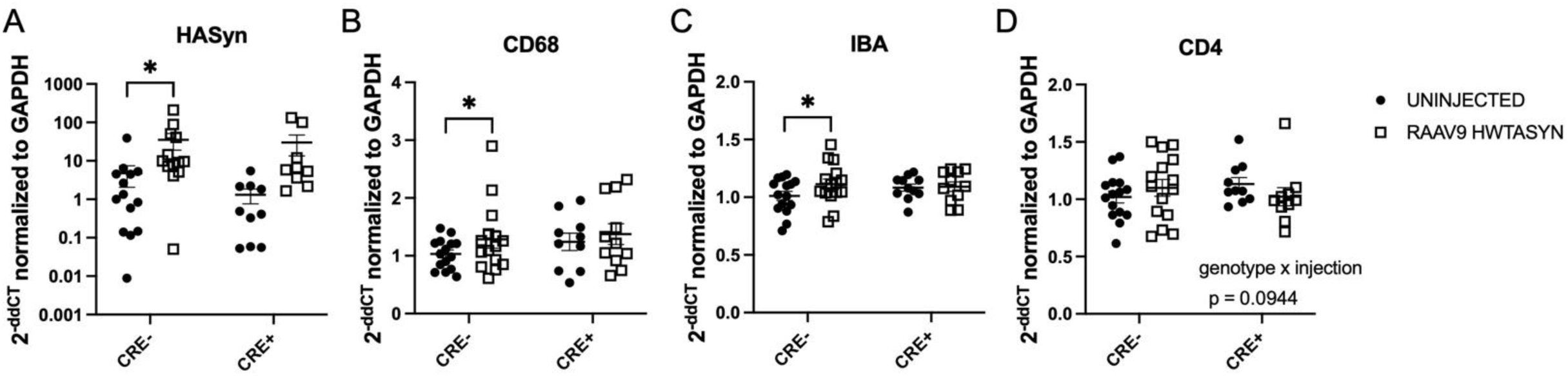
Inflammatory responses in the striatum of MHCII-deficient mice 4 months after nigral rAAV9 hWT-ASYN injection is significantly dampened. RNA extracted from microdissected striatum from the uninjected hemisphere and rAAV2/9 human WT α-synuclein (RAAV9 hWT-ASYN) injected hemisphere of LysMCre-I-Abfl/fl (CRE-) and LysMCre+I-Abfl/fl (CRE+) mice. Gene expression fold change was quantified using the double delta CT method relative to GAPDH and CRE-group mean with RT-PCR. (A) CRE-animals demonstrated increased levels of human α-synuclein transcript in the hemisphere ipsilateral to the injection compared to the contralateral hemisphere. Similarly, CRE-injected hemispheres had elevated levels of (B) CD68 and (C) IBA1 compared to the uninjected hemisphere. (D) A trend was found for a genotype and injection interaction when evaluating CD4 gene expression (p=0.0944). RT-PCR triplicates that had standard deviations higher than 0.3 were excluded. Data are plotted as mean ± standard error of the mean. P values are indicated for two-way repeated measures ANOVA and an uncorrected Fisher’s LSD multiple comparison (mixed effects model was used for analysis using missing values due to standard deviation exclusion criteria): *p<0.05.

### The high-risk HLA-DRA rs31298882 GG genotype is associated with increased inducibility of MHCII proteins in cryo-recovered PBMCs

To confirm previous findings that the high-risk SNP genotype GG at rs3129882 is associated with greater MHCII inducibility, cryopreserved PBMCs were treated with IFN-γ and HLA-DR and -DQ alpha and beta chain gene expression was measured. Analysis correcting for multiple comparisons found the greatest IFN-γ-induced upregulation of HLA loci in PBMCs from control *AA* (CAA) and PD *GG* (Figure 7). Total PBMCs from PD *GG* demonstrated significant inducibility of HLA-DRB compared to PD *AA* when stimulated with IFN-γ, but this effect was not seen in healthy control PBMCs of similar genetic polymorphism (Figure 7B). These data are consistent with findings reported in previous work done on fresh PBMCs, suggesting cryopreservation does not compromise the SNP or disease impacts on HLA expression^30^.

**Figure 7.**
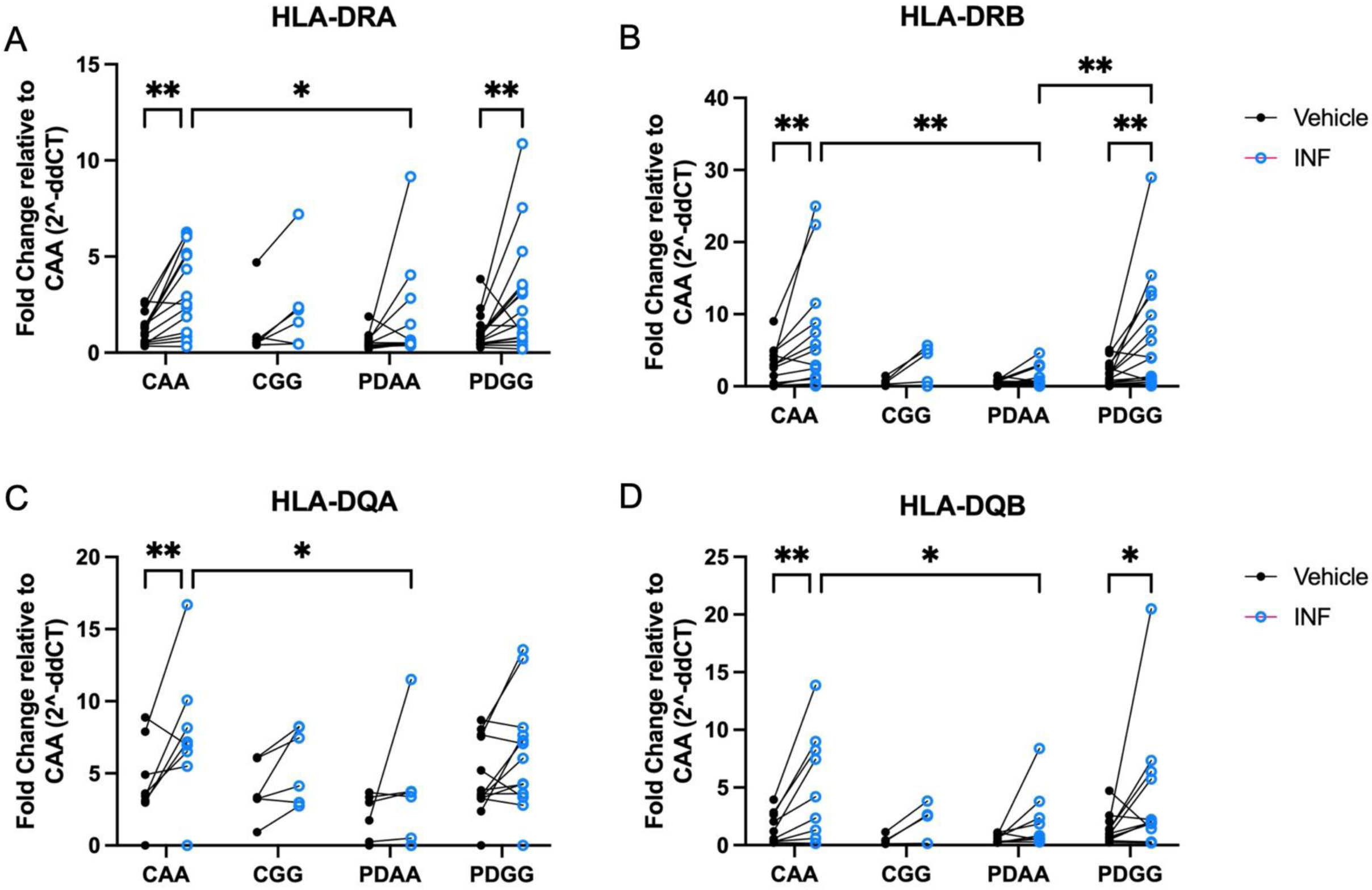
The high-risk HLA-DRA rs31298882 GG genotype is associated with increased inducibility of MHCII proteins in cryorecovered PBMCs. Individuals with PD with the high-risk *rs3129882* GG genotype show greater IFN-γ inducibility compared to AA individuals. HLA-DR and -DQ alpha and beta chain gene expression was measured from cryopreserved PBMCs following thawing and stimulation with vehicle and IFN-γ. Gene expression fold change was quantified using the double delta CT method relative to GAPDH and control AA group with RT-PCR. Inducibility from IFN-γ when compared to vehicle treatment was found in (A) HLA-DRA, (B) HLA-DRB, (D) HLA-DQB in control AA (CAA) and Parkinson’s disease GG (PDGG), (C) yet this effect was only seen in CAA samples when evaluating HLA-DQA. (B) HLA-DRB inducibility was significantly increased in PDGG compared to PDAA individuals. A two-way ANOVA with uncorrected Fisher’s LSD multiple comparisons was used to evaluate HLA expression between groups (control/PD with either AA or GG) and cell treatment (vehicle vs IFN-γ). *p<0.05, **p<0.01.

### Flow cytometry confirms genotype- and disease-specific effects of HLA rs3129882 SNP detected in peripheral blood from individuals with Parkinson’s disease and controls

To assess the effect of the high-risk rs3129882 SNP on T-cell subset frequency, PBMCs from PD and healthy control subjects were prepared for flow cytometry. Due do the SNP genotype differences we observed in MHCII mRNA inducibility, we hypothesized that there would be alterations in CD4+ T cell subsets in the *GG* group relative to the *AA* group. We observed no significant differences in overall frequencies of total CD3+ T cells among our four groups (Figure 8A). No significant differences were seen between genotypes for Treg, Th17, or CD4:CD8 ratio (Figure 8B-D). Flow analysis revealed a trend toward reduced total CD8+ T cells in subjects with *GG* genotype (p=0.0514; Figure 8E), while evaluation of CD8 subsets showed a significant SNIP effect in CD8 central memory cells, decreased CD8 naïve cells in PD *AA* participants relative to health control *AA* participants, and increased CD8 effector T cells in PD *AA* relative to healthy control *AA* participants (Figure 8F-I). We observed a statistically significant higher frequency of CD4+ T cells (CD3+CD4+) in *GG* participants where multiple comparisons revealed both PD and healthy controls with GG had elevated CD4+ cell frequency compared to PD *AA* genotype (Figure 8J). Additionally, CD4+ effectors (CD45RA+/CCR7-) were significantly increased in PD *AA* compared to control *AA*, yet no statistically significant differences were found in other CD4+ T cell subsets frequencies (central memory, effector memory, naïve; Figure 8K-N) among our four groups. Evaluation of PBMC counts (cells/μl) show similar patterns of change for T-cell effector subsets (Supplementary Figure S3).

**Figure 8.**
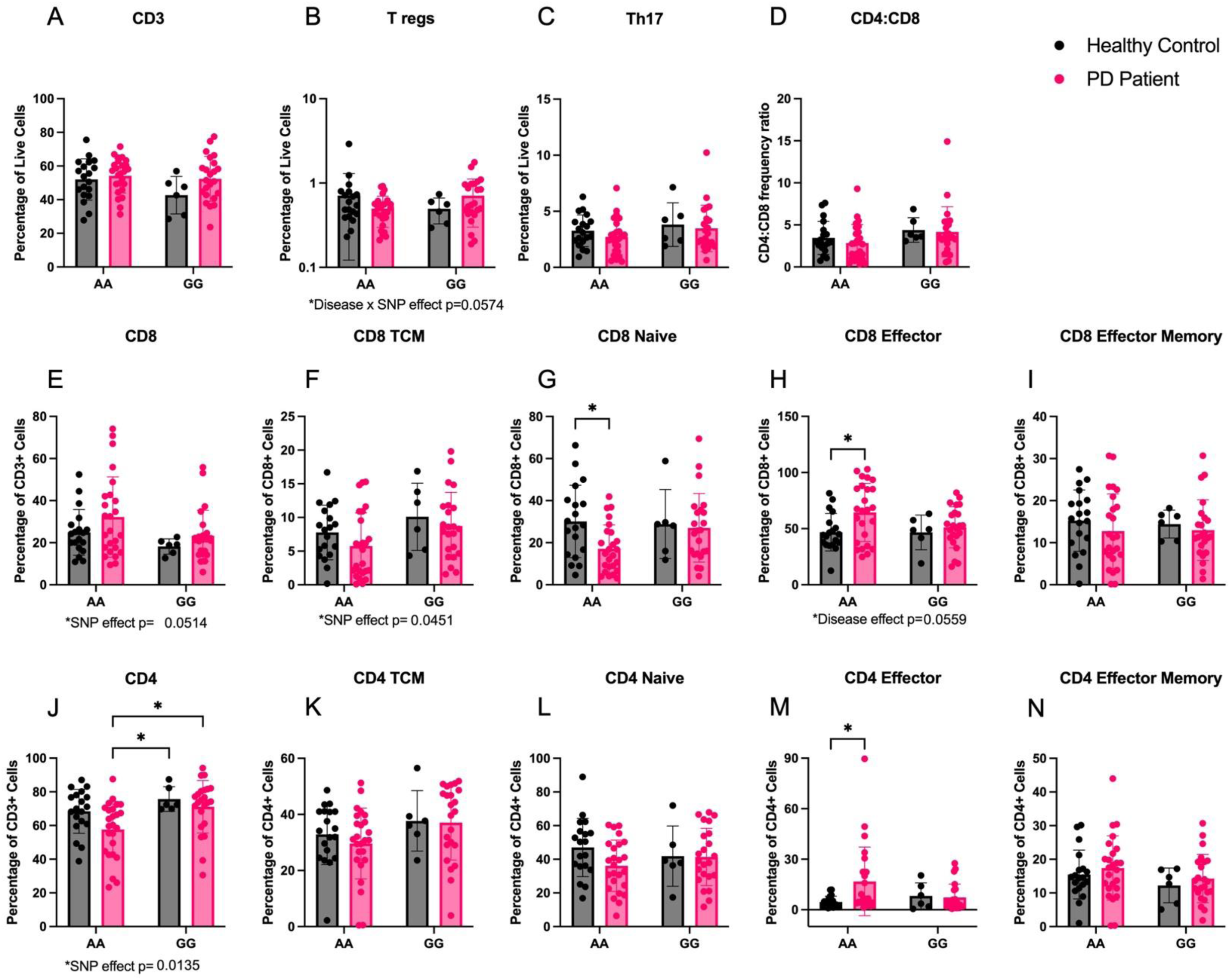
The high-risk GG genotype rs3129882 is associated with increased frequency of CD4+ independent of disease while the low-risk AA genotype is associated with decreased naïve CD8 and increased CD8 effector T cells but only in individuals with PD Flow cytometry staining was used to determine frequencies of T cells in cryopreserved PBMC samples from healthy controls (HC) and individuals with PD with the *rs3129882* genotype AA or GG (increased risk of PD). HC AA n=19, HC GG n=6, PD AA n=25, PD GG n=23. (A-C) No effect of disease or genotype was identified in frequency of CD3+ T cells, T regulator cells (CD3+CD4+CD25+CCR6+), or TH17 cells (CD3+CD4+CD25+CCR6+ CCR4+) out of total live cells. (D) Ratio of CD4:CD8 cell frequency demonstrates no differences between groups. (E,F) An effect of SNP genotype was seen in the frequency of total CD8+ T cells out of CD3+ cells (p=0.0514) and CD8+ central memory (CD8+CCR7+CD45RA-) out of CD8+ cells (p=0.0451). (G-I) CD8+ cell subtype frequencies were evaluated and CD8+ naïve (CD8+CCR7+CD45RA+) and effector cells (CD8+CCR7-CD45RA+) were found different between disease when expressing the AA genotype, while no effect was found in effector memory cells (CD8+CCR7-CD45RA-). (J) Frequency of CD4+ cells out of CD3+ cells are significantly higher in those with the GG genotype compared to PD AA. (K-L) Evaluation of CD4+ cell subtype frequencies found no differences in central memory (CD4+CCR7+CD45RA-), naïve (CD4+CCR7+CD45RA+) or effector memory (CD4+CCR7-CD45RA-), but increase effector cells (CD4+CCR7-CD45RA+) in PD AA compared to HC AA. Two-way ordinary ANOVA with Tukey correction for multiple comparisons was used to test for significant differences. Data are plotted as mean ± standard error of the mean. Two-way ordinary ANOVA with Tukey correction for multiple comparisons used to analyze the data. Trending or significant disease, SNP, and interaction effects noted below each graph. *p<0.05

To understand how SNP genotype and disease state may impact the relationship between MHCII gene expression and T cell populations, we performed a correlational analysis of baseline gene expression of HLA-DR and DQ alpha and beta chain and T cell populations identified through flow cytometry from samples belonging to the same patient. From these analyses, we identified 5 significant correlations in our PD *GG* group, most of which related to HLA-DRB gene expression. Here we find that PD *GG* individuals exhibit a positive relationship between HLA-DRB gene expression and CD4 naïve T cells frequency, while also exhibiting an inverse relationship with CD4 central memory cell frequency, T regulatory (Treg) cell frequency, and total CD8 T cell frequency (Figure 9A-D). Moreover, we find the HLA-DQB in PD *GG* displays a positive relationship to naïve CD4 T cell frequency as well (Figure 9E). Overall, these data demonstrate that increased baseline expression of HLA in PD GG positively correlates with naïve CD4 T cell populations but negatively correlates with effector T cell subsets.

**Figure 9.**
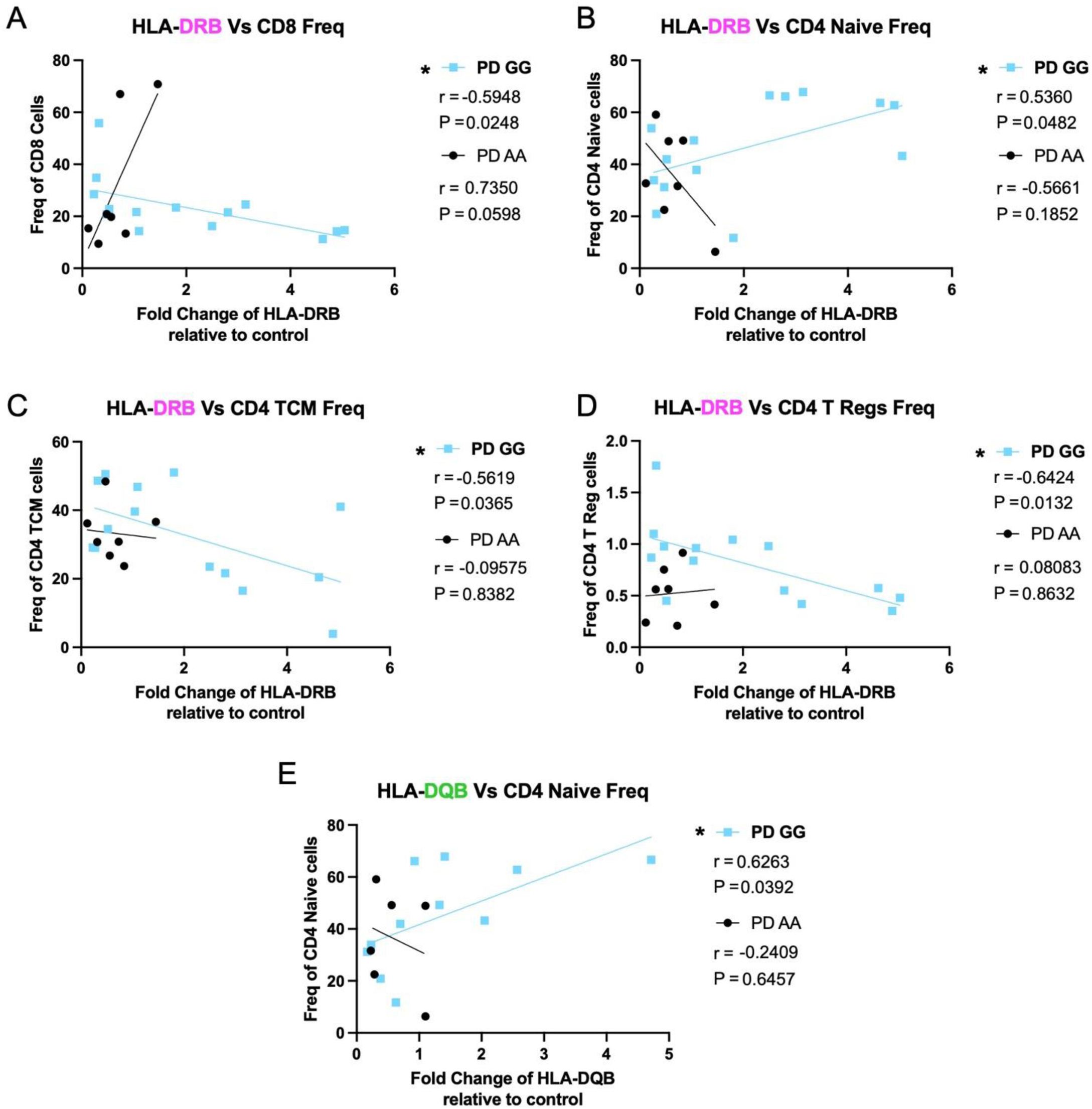
HLA gene expression positively correlates with naïve CD4 T cell populations but negatively correlates with effector T cell subsets in PD GG. Baseline fold change in HLA gene expression relative to GAPDH correlates to frequency of T cell subsets in individuals with PD with GG SNP genotype at *rs3129882* compared to PD AA. PD AA n= 6-7, PD GG n =11. (A) HLA-DRB gene expression correlated with frequency of CD8+, (B) CD4+ naïve cells, (C) CD4+ Central Memory cells, and (D) CD4+ Tregs. (E) HLA-DQB gene expression correlated with frequency of CD4+ naïve cells. Samples were analyzed with two tailed Pearson’s Correlation test with 95% confidence interval. R and P values of correlations present under their respective PD SNP group. * p<0.05.

### ATAC-seq analysis of the HLA locus reveals genotype-as well as disease-dependent differentially accessible chromatin regions in human monocytes

To further understand the if the SNP genotype was associated with global or local epigenetic changes, we profiled the chromatin accessibility landscape of IFN-γ stimulated and unstimulated monocytes with from HC and PD patients harboring the GG and AA alleles. A principal component analysis (PCA) of all accessible regions across the experimental cohort revealed a major change in accessibility following IFN-γ stimulation in all samples (Figure 10A). PCA analysis did not reveal any obvious variation in samples based on SNP genotype or HC versus PD status. The rs3129882 SNP is located in the first intron of the *HLA-DRA* gene in a region that harbors minimal accessibility (Figure 10B). Therefore, to determine of other accessible regions of the genome were altered, we compared all sample groups pairwise. We identified very few significant differentially accessibility chromatin regions (DAR) when looking at the data globally across patient groups or by genotype, yet 10-13k pairwise DAR were identified in response to IFN-γ, which served as a positive control (Figure 10C). We next focused our analysis on 25 accessible regions within the MHCII locus that covered known regulatory regions with high variability across the human population including a superenhancer (DR/DQ-SE)^38^ and the CTCF bound insulator elements that surround the *HLA-DR* and *HLA-DQ loci (C1, C2*, and *XL9*)^38–40^. The superenhancer and CTCF sites are linked by chromatin looping interactions that are required to fully express the locus. This focused analysis found 3 conditions where differences were observed, including one peak that had higher accessibility in both HC and PD GG genotypes compared to PD AA in response to IFN-γ (Figure 10D). The other region was more highly accessible in PD AA compared to PD GG in naïve monocytes (Figure 10D). Each of the DARs within the HLA locus were visualized using a genome plot with comparisons across all groups to highlight the specificity of the changes at these regions compared to the other sites within the greater MHCII locus (Figure 10E-F).

**Figure 10.**
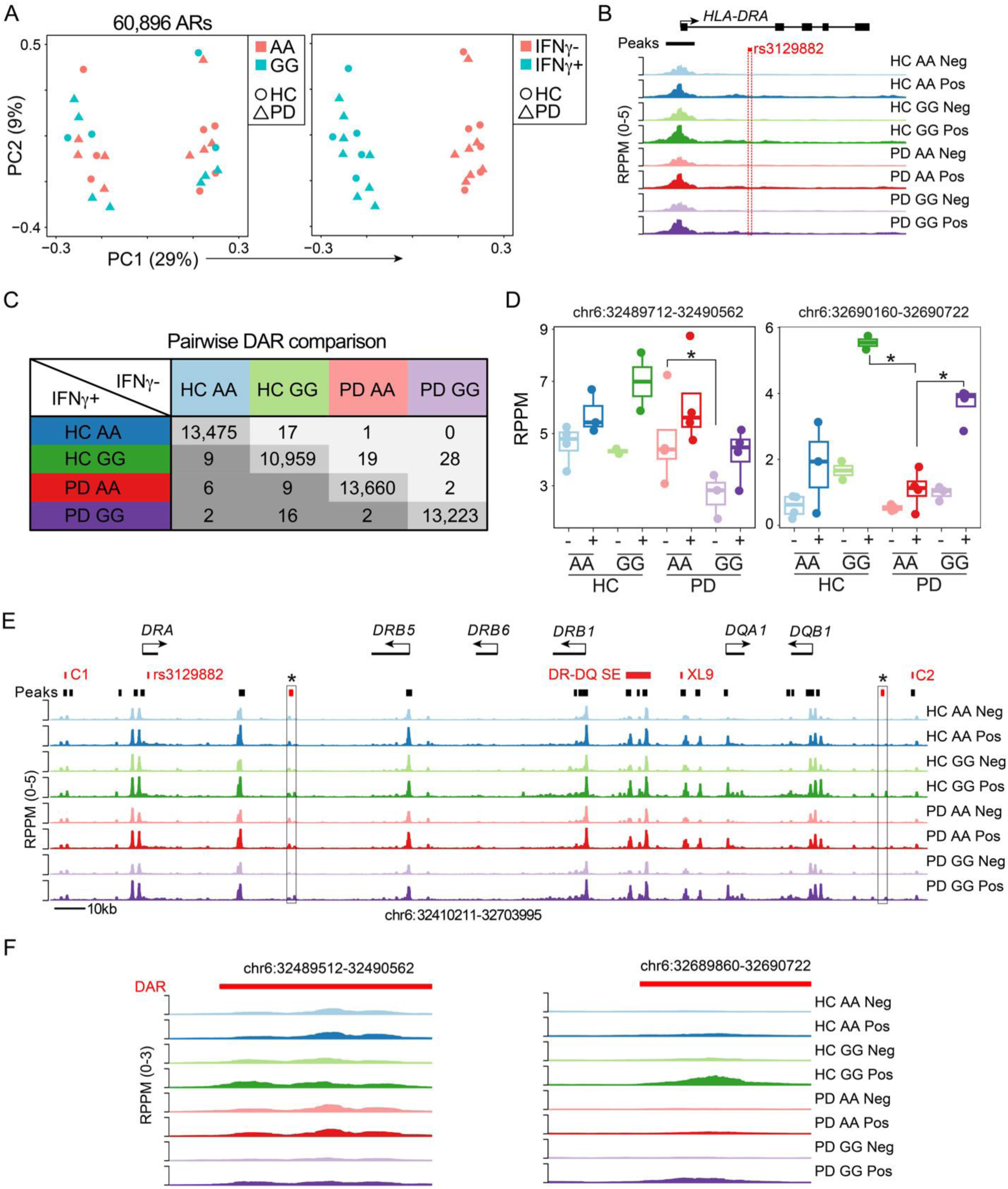
ATAC-seq analysis of the *HLA* locus reveals genotype as well as disease-dependent differentially accessible chromatin regions in human monocytes. To determine if carriers of the homozygous AA or GG rs3129882 genotype and/or PD resulted in epigenetic changes in the MHCII locus, monocytes were purified and treated with control or 100U of IFN-γ for 18hr and ATAC-seq performed to measure global changes in chromatin accessibility. (A) Principal component analysis (PCA) of 60,896 accessible regions (AR) showing the distribution of healthy donors (HD, circles) or Parkinson’s Disease (PD, triangles) monocytes. The samples are color coded by genotype (left) or IFN-γ treatment (right). (B) Genome plot of the *HLA-DRA* locus showing the average reads per peak per million (RPPM) normalized ATAC-seq signal in each group. The location of an accessible peak at the promoter is indicated in black and rs3129882 SNP in red. (C) Table of differentially accessible chromatin regions (DAR; absolute log_2_FC >1 and FDR < 0.05) for all pairwise comparisons. Note along the diagonal large numbers of DAR in response to the IFN-γ stimulation compared to the small number of DAR between both disease status and rs3129882 genotypes. (D) For the 25 accessible regions within the MHCII locus between the C1 and C2 CTCF sites a focused differential analysis revealed three DAR. Box plots showing the accessibility of each group across the two DAR. (E) Genome plot depicting the MHCII locus denoting the location of genes and accessibility in each condition. In red are major elements of the locus including the CTCF sites (C1, XL9, C2), Super Enhancer (DR-DQ SE), and rs3129882 SNP. Black bars indicate the location of accessible regions with the two DAR indicated by a red box and an asterix. (F) Genome plots showing the accessibility patterns for the two DAR identified in D. * indicates that comparison was significant.

## Discussion

The work herein was motivated by a seminal finding that a common variation in the gene for MHCII is associated with increased risk for late-onset PD which our group demonstrated has functional consequences for individuals with the high-risk *GG* genotype at the HLA-DRA SNP *rs3129882.* Specifically, these individuals displayed monocytes in blood with higher baseline MHCII gene and protein expression as well hyper-responsiveness to an immune challenge in conjunction with PD status^30^. Therefore, we set out to investigate the role of MHCII protein expression in a mouse model of PD-like pathology. Because this mouse displays normal CD4+ and CD8+ T cell frequencies in PBMCs, it enabled us to more directly draw conclusions about the role of MHCII-based antigen presentation, rather than in the general absence of a functional adaptive immune system. We originally set out to reduce MHCII peripherally to understand the differential contribution of peripheral versus central antigen presenting immune cells to alpha-synuclein-induced inflammation and neurodegeneration. The choice to drive Cre expression under the LysM promoter was based on original papers that described LysMCre-mediated deletion as specifically targeting peripheral myeloid populations such as macrophages and granulocytes^41^. Because more recent studies have highlighted the possibility of lysozyme upregulation under inflammatory conditions in microglia^42^, it is not surprising that we ultimately observed a trend toward reduced frequency of MHCII+ microglia (p=0.0638, Figure 2) and significant reduction of MHCII MFI on microglia in mice overexpressing human alpha-synuclein. The data in Figure 2 suggest that microglia and other myeloid cells within the brain have different levels of LysM expression, as shown by the mild effects on frequency of MHCII+ microglia compared to more significant reduction in frequency of MHCII+ monocytes and granulocytes.

Our results demonstrate that the reduction of MHCII from LysM-expressing myeloid cells produces mild protection of the striatal dopaminergic phenotype, which may be attributed to the globally altered T cell subsets and dampened inflammatory responses in the striatum. Given the T cell subset changes in mice with reduction of MHCII, we expected to see inverse T cell subset changes in humans with increased MHCII inducibility. We expected that more MHCII would be associated with higher frequency of pro-inflammatory T cell subsets. However, in a cohort of individuals with PD harboring the high-risk *rs3129882 GG* genotype, the expected expansion of T cell effector cell populations was not observed despite MHCII inducibility in bulk PBMCs. Instead, the *GG* population exhibited increased *naïve* CD4+ T cells, which may be unrelated to SNP genotype and MHCII expression or related to PD. Another possibility is that the MHCII expressed in *GG* individuals is, for unknown reasons, incapable of inducing appropriate CD4+ cell differentiation. Future work should investigate structural and functional differences in the MHCII protein in individuals with the high-risk *rs3129882 GG* genotype. Despite these surprising findings related to MHCII function, human monocytes differentially accessible chromatin regions were identified in the MHCII locus in both a disease- and genotype-dependent manner. Overall, these results support a critical role for MHCII in α-synuclein-induced immunogenicity.

Several previous investigations, were not designed to distinguish different contributions to antigen presentation by peripheral versus brain-resident myeloid immune cells. Until more recent work using CX3CR1- and TMEM119-driven Cre expression for targeted, cell-specific of MHCII, it had been impossible to distinguish between peripheral and CNS-resident macrophage antigen presentation contributions to neuroinflammation and α-synuclein-induced neurodegeneration^25^. The distinction is of critical importance to translational work, as we must consider the challenges of developing therapeutics to prevent or treat PD that may need to penetrate the blood-brain barrier. Recent work that elegantly and meticulously differentiated between microglia, BAMs, and peripheral innate immune cells suggests that therapeutic interventions likely need only engage BAMs in order to have a therapeutic benefit, as it is the BAM population that is actively engaged in the antigen presentation driving inflammatory cytokine release during α-synuclein-related degeneration^25^. While not cell-type specific, our work reinforces the conclusion that MHCII expression is essential for α-synuclein-related degeneration – the degeneration is modified, though not entirely absent, in the CRE+ mice which inherently display decreased myeloid MHCII expression. LysM (also known as Lyz2) is known to be expressed in BAMs^43^, suggesting MHCII reduction in BAMs in our CRE+ mice; however, this specific cell-type was not evaluated in this study.

Mechanistically, based on the data reported by our group^30^, we hypothesized that decreased MHCII expression on any innate immune cell population would benefit the health of dopaminergic neurons via decreases in effector CD4+ T cell frequency; and it was of importance to determine whether Treg cells, which suppress inflammation^44^ and can attenuate α-synuclein-induced neurodegeneration with early expansion^45^, were affected or not. We observed a decrease in the frequency of PBMC and DCLN Tregs in CRE+ mice, suggesting inflammatory immune responses in CRE+ mice may persist after an insult relative to CRE-controls. Although DCLN contain lymphocytes draining from the brain, we did not directly evaluate Tregs in the brain. Furthermore, this study did not evaluate Treg populations throughout the post-α-synuclein injection period but only at endpoint, and therefore we cannot assess the effect that altered Treg frequency had on the acute inflammatory response to human α-synuclein in the SN. Future work should assess the presence of Tregs in the brain by flow cytometry at time points closer to versus farther away from the onset of α-synuclein expression. In addition to the shift in Treg frequency, CRE+ mice have decreased frequencies of central memory and effector memory CD4+ T cells and an increase in the frequency of naïve CD4+ T cells. Interestingly, individuals with PD have been reported to have increased frequency of a marker of memory subsets CD45RO+ T cells^46^. If we assume the T cell frequencies of an individual with PD are disease-promoting or at least permissive to progression of disease, our data indicate that the reduction of MHCII on peripheral myeloid cells is beneficial, in that it promotes the development of a T cell compartment lacking the characteristics seen in individuals with PD. Our results revealed that, similar to PBMCs, the frequency of central memory CD4+ T cells in the brain and effector memory CD4+ T cells in mouse DCLN was decreased in CRE+ mice. These observations are consistent with our finding that MHCII is reduced on peripheral myeloid cells and provide confirmation that the targeted reduction in LysM*-*expressing cells is sufficient to affect CD4+ T cells populations globally. If achieved in individuals with PD, decreased CD4+ T cell traffic to the brain could protect dopaminergic neurons^35^ from an adaptive immune response directed against α-synuclein or other immunogenic neuron-derived antigen(s).

In previous work from our lab, it was observed that the high-risk *GG* genotype at the *rs3129882* SNP in the HLA-DRA gene synergizes with exposure to pyrethroid pesticides to increase risk for PD^30^. As our mouse data suggests that loss of MHCII may modify or potentially mitigate neurodegeneration via shifts in the CD4 T cell repertoire, we were surprised to see that individuals with PD with the high-risk *rs3129882 GG* genotype display a positive correlation between increased HLA expression and naïve T cell populations. A potential interpretation of this observation is that increased HLA-DRB expression is a compensatory mechanism for MHCII protein with *decreased* antigen presentation capacity in individuals harboring *GG* and having PD. The increased inducibility of HLA does not result in the expansion of effector cell populations and the decrease of naïve populations as we suspected; rather, the opposite was observed. It is also a possibility that variation in disease duration within our PD cohort may be a variable impacting our results and/or interpretation of the findings. Previous work has suggested that shorter disease duration is associated with more pro-inflammatory cytokine secretion, followed by a more Th2/Th17-focused response that emerges later in disease. The connection between MHCII expression and activation and differentiation of CD4+ helper T-cell subsets may fluctuate in PD *GG* individuals over time. Further experiments to investigate the functionality of MHCII in those with the *GG* genotype will need to be conducted to explore these possibilities, with particular attention to the capacity for expression of α-synuclein-derived peptides, as it is known that individuals with PD possess α-synuclein-specific T cells^20^.

As the *rs3129882* SNP is located in the first intron of the *HLA-DRA* gene, we tested the hypothesis that it was in linkage disequilibrium with epigenetic changes within the MHCII locus that could result in altered HLA-DR expression. Using chromatin accessibility for enhancer activity, an analysis spanning disease and genotype revealed few global differences, likely indicating the effect is confined within the MHCII locus. The focused analysis of peaks identified two regions with altered accessibility and one that was elevated in both HD and PD GG monocytes treated with IFN-γ. Through previous studies of the MHCII superenhancer (DR/DQ-SE) we showed that deletion of the super enhancer regulatory elements led to a two-fold reduction in HLA-DR expression and an altered ability to activate CD4 T cells^38^. This indicates that even subtle changes in HLA surface expression can have important consequences on the ability to induce immune responses. Additionally, it is notable that the *rs3129882* SNP itself was not associated with significant changes in chromatin accessibility suggesting that it does not represent what would be expected as a binding site for a transcription factor. Other studies localizing PD risk SNPs to open chromatin of microglia indicate that altered cis-regulatory element activity also has a role in the etiology of PD^47^.

Taken together, our work reinforces the conclusion that MHCII expression is essential for α-synuclein-related degeneration and inflammation, and that changes in MHCII expression and inducibility mediated by the high-risk GG SNP not only influence human T cell frequencies but also determines chromatin accessibility in a disease- and genotype-specific manner. These findings have implications for clinical trial recruitment and stratification, as well as for incorporation of functional measures of innate and adaptive immune dysfunction in the peripheral blood of at-risk individuals and people living with Parkinson’s into trials to monitor responsiveness to immunomodulatory interventions.

## Methods

### Animals

Two lines of mice were purchased from Jackson Labs: LysMCre mice B6N.129P2(B6)-Lyz2^tm1(cre)Ifo^/J (Jackson Labs catalog # 018956) and I-Ab-flox mice B6.129X1-H2-Ab1^b-tm1Koni^/J (Jackson Labs catalog # 013181) were crossed to generate I-Ab-deficient (LysMCre+I-Ab^fl/fl^ or CRE+) and control (LysMCre**-**I-Ab^fl/fl^ or CRE-) male mice for this study. Mice were housed in the facilities in the Whitehead Biomedical Research building in the Division of Animal Resources at Emory University with *ad libitum* access to food and water. The animal facility is on a 12-hour light/dark cycle. Experimental procedures were approved by the Institutional Animal Care and Use Committee of Emory University and all experiments were performed in accordance with relevant guidelines and regulations and in compliance with the National Institutes of Health guide for the care and use of laboratory animals. Furthermore, the study is reported in accordance with ARRIVE 2.0 Essential 10 guidelines. Mice were divided into 2 cohorts and followed an experimental design outlined in Figure 1. Cohorts 1 and 2 consisted of mice that received stereotaxic injection at 2-3 months old (young adult). Cohort 1 (n= 11 CRE+ and n=16 CRE-) was euthanized four months post-injection by intraperitoneal Euthasol (Virbac Animal Health, Fort Worth, TX) and saline perfusion for PBMC and DCLN flow cytometry as well as Western blot, qPCR, and immunohistochemistry. Cohort 2 (n=5 CRE- and n=6 CRE+) was euthanized four months post-injection via isoflurane-anesthetized cervical dislocation and non-perfused brains immediately collected for brain immune cell flow cytometry.

### Stereotaxic Injections

The rAAV9 hWT-ASYN was generously provided by Dr. Frederic Manfredsson (Stock Titer: 2.1×10^12^)^48,49^. Mice were unilaterally injected (2μL) under stereotaxic guidance to the left SN as in^4^. Surgery was performed under 1-2% isoflurane anesthesia. Mice were placed into stereotaxic frame and injection coordinates for the SN were determined relative to bregma: AP -3.2mm, ML +1.2mm, DV -4.6mm from dura. The virus was delivered via handmade glass microcapillary pipette affixed to an 80000 Hamilton syringe (Hamilton Company) coated in Sigmacote (Millipore Sigma Cat#SL2). Injections using a glass microcapillary tip have previously shown to reduce immunodetection of IgG in the parenchyma compared to the use of Hamilton syringes/needles, suggesting that the use of a glass tip to deliver AAV mitigates BBB breakdown^50^. Post-operative pain relief was provided using buprenorphine (Reckitt Benckiser, NDC 12496-0757-1, 0.1mg/kg, intraperitoneal).

### Tissue collection

In order to assess the extent of MHCII reduction in the peripheral myeloid cells of CRE+ relative to CRE-mice, splenocytes, peritoneal macrophages were harvested from naïve animals (did not receive virus) and prepared for flow cytometry. Spleens were placed in 1xHBSS and passed through a 40μm cell strainer to create a single cell suspension. Following centrifugation and removal of supernatant, red blood cells were lysed using ACK buffer. RPMI media was used to wash away ACK buffer, and splenocytes were counted using a hemacytometer and Trypan blue exclusion. Splenocytes were loaded into the autoMACS Pro Separator (Miltenyi Biotec, Bergisch Gladbach, Germany) for CD11b+ cell selection. Peritoneal macrophages were harvested by following a previously published protocol^51^. Briefly, mice were administered 3% Brewer thioglycolate medium into the peritoneal cavity and 72 hours later a peritoneal lavage was performed using an 18g needle and 5mL of RPMI 1640. Cells were recovered, counted, and went through the autoMACS CD11b+ isolation described for spleen. Cells were stained for flow cytometry (see Table 1 for list of antibodies) and populations of splenocytes, and peritoneal macrophage defined by specific gating strategies (Supplementary Figure S4 and S5, respectively).

**Table 1.**
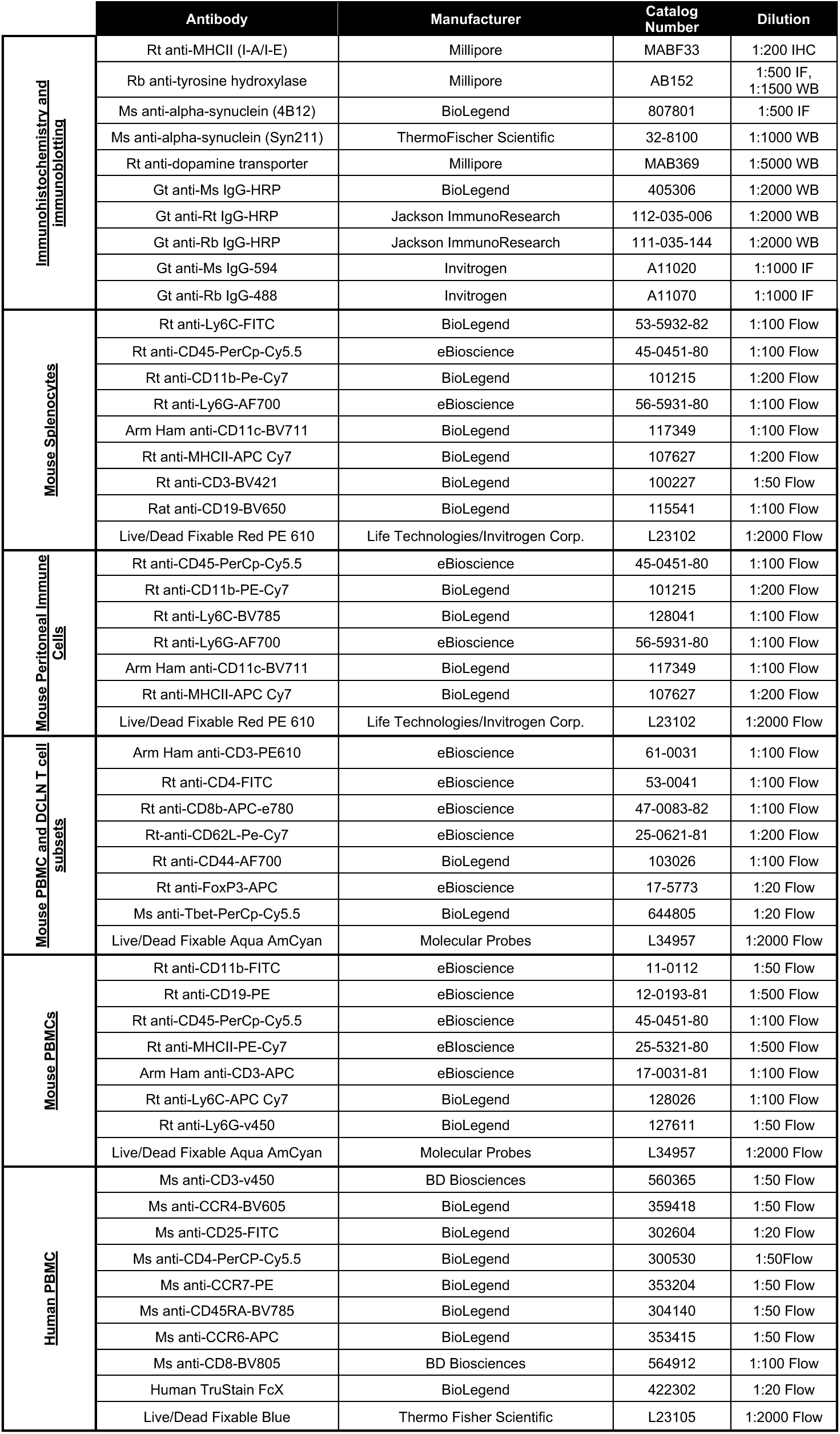

For cohort 1 (PBMC and DCLN flow cytometry studies, striatal protein and RNA assessments, and immunohistochemistry), mice were injected with Euthasol followed by saline perfusion for five minutes. Brains were extracted and the left and right striatum were dissected, and flash frozen in liquid nitrogen. Tissue was stored at -80°C until processed for Western blot. The caudle portion of the brain (posterior of striatum) was placed in 4% paraformaldehyde at 4°C for 24 hours. Tissue was transferred to a 30% sucrose solution in phosphate buffered saline (PBS) and stored until sliced. Sectioning was performed on a freezing stage microtome (Leica SM 2010R, Leica Biosystems Inc., Buffalo Grove IL) and 40μm coronal slices were stored in PBS with 0.1% sodium azide at 4°C until stained.

Approximately 200uL of whole blood was collected from cheek bleeds prior to Euthasol injection. Blood was collected in EDTA coated tubes (Coidien Cat#8881311248) and 100uL was treated with 1x red blood cell lysis buffer (Biolegend Cat#420301) in order to isolate PBMCs. PBMCs were divided into two wells. One well was stained using the mouse immunophenotyping panel and the other was stained using the mouse T cell subset panel (Supplementary Figure S6, see below for flow cytometry methods). The right and left DCLNs were removed from each mouse and placed into 1xHanks Balanced Salt Solution with 5% fetal bovine serum (FBS). A single cell suspension was made by grinding the tissue through a 70μm filter. All cells isolated from DCLNs were plated and stained for multi-color flow cytometry using the mouse T cell subset panel (Supplementary Figure S6, see below for flow cytometry methods).

For cohort 2 (brain flow cytometry), mice were isoflurane-anesthetized for cervical dislocation and brains were collected. The caudal portion of the brain, including much of the hippocampus and midbrain, was prepared for immune cell isolation and flow cytometry as previously described by our group with some modifications^52,53^. The caudal brain was minced in 2mL of RPMI 1640. Samples were placed in 2.5mL of dissociation medium (1.4U/mL collagenase VIII [Sigma-Aldrich Cat#C1889], 1mg/mL DNase1 [Sigma-Aldrich Cat#DN25] in RPMI1640) and incubated at 37*°*C, 5% CO_2_ for 15 minutes, inverting every 5 minutes. Samples were then neutralized with 5mL of 10% FBS (Atlanta Biological) in RPMI1640 warmed to 37*°*C. Tissue was pelleted and the supernatant was aspirated. The pellet was then homogenized in 3mL of 1xHanks Balanced Salt Solution (HBSS) using fire-polished glass pipettes. Dissociated cells were filtered through a 70μm strainer with 5mL of 1xHBSS and then pelleted. At room temperature, the pellet was resuspended in 4mL of 37% Percoll (Percoll pH 8.5–9.5; Sigma-Aldrich, Cat#P1644), with 70% Percoll layered below, 30% Percoll layered above, and 1xHBSS on top. The Percoll gradient was centrifuged, and brain immune cells were collected from the interface between the 70% and 37% Percoll layers. Cells were washed with 1xHBSS and stained for flow cytometry and populations defined with a gating strategy to identify microglia, monocytes and lymphocytes (Supplementary Figure S7). Gating was based on strategies previously published by our group^52^.

### Mouse flow cytometry

For all panels, cells were incubated with Live/Dead Fixable (Table 1) and anti-mouse Fc block (CD16/CD32) (1:100, eBioscience Cat#14-0161085) along with cell surface markers in PBS. Following incubation with surface markers, intracellular staining was performed for the mouse T cell subset panel samples. To fix and permeabilize, the Invitrogen intracellular staining kit was used as per the manufacturer’s protocol (eBioscience Cat#00-5523-00). Spherobeads (Spherotech Cat#SRCP-35-2A) and OneComp beads (Invitrogen Cat#01-1111-42) were used to set voltages on the cytometer and compensation settings, respectively. Samples were immediately run on an LSRII (BD Bioscience) following staining by an experimenter blinded to the treatment groups. FlowJo (version 10) was used for analysis.

### Immunohistochemistry

From Cohort 1, nigral slices were stained for MHCII analysis using previously published protocol^54^. Briefly, endogenous peroxidase was blocked by a 15-minute incubation in 3% hydrogen peroxide. Tissue was washed and then incubated in blocking solution consisting of 10μg/mL avidin, 8% normal serum and 0.1% triton-x for 1 hour at 4°C and followed by an overnight incubation in 50μg/mL biotin, 2% normal serum and primary antibodies specific for MHCII (Millipore MABF33, room temperature incubation). Tissue was further covered in biotinylated secondary antibodies with 2% normal serum for 1 hour at 4°C and amplified with ABC (Vectastain Cat#PK-6100) and developed with diaminobenzidine tablets. Tissue was mounted on slides, dehydrated in graded ethanols and cleared with xylene before adhering coverslip with cytoseal mounting media.

### Western blots

From cohort 1, striatal tissue was homogenized using a pestle motor mixer in RIPA buffer with protease and phosphatase inhibitors. Following a bicinchoninic acid protein assay, samples were diluted in water and 2x Laemmli buffer. Samples were boiled for 5 minutes and run immediately for Western blot. Pre-cast, 15-well 10% mini-PROTEANâ TGX^TM^ polyacrylamide gels from Bio-Rad were loaded with 10μg of protein from each sample. Gel electrophoresis was performed at 100V for 110 minutes. Methanol-activated PVDF membranes were used for protein transfer via the Trans-Blot Turbo System (Bio-Rad). The high molecular weight, 10 minutes setting was used to transfer protein. REVERT stain (LI-COR Biosciences Cat#926-11011) for detection of total protein was performed and membranes were imaged in the 700 channel with a 30 second exposure. The membrane was then blocked in 5% milk in 0.1% Tween-20 TBS buffer for 1 hour at room temperature. Blots were incubated overnight at 4°C in 5% milk buffer with primary antibodies against mouse dopamine transporter (DAT), and TH at the concentrations indicated in Table 1. Blots were then washed and incubated with secondary HRP conjugated antibodies (1:2000) at room temperature for 1 hour. Following additional washes, blots were developed using West Pico Chemiluminescent Substrate or SuperSignal West Femto Chemiluminescent Substrate (ThermoFischer Scientific Cat#34096). Band intensity was measured using an Azure c400 (Azure Biosystems) and analysis was performed using ImageStudio Software (Li-Cor Biosciences). Samples were normalized to their own total protein bands and relative expression determined as compared to the average values from the uninjected hemisphere of Cre-animals.

### Mouse quantitative real-time polymerase chain reaction (RT-PCR)

To complement our understanding of protein level changes to the nigrostriatal DA system following viral vector-mediated expression of human α-synuclein, RNA was isolated from the same left and right striatal samples used for Western blots from cohort 1. RNA was isolated using TRIzol^TM^ reagent according to the manufacturer’s protocol (ThermoFischer Scientific Cat#15596018). Reverse transcription was performed to make cDNA as published^30^. RT-PCR was performed to detect transcript levels of human α-synuclein, surface makers of immune cells (CD4, MHCII, MHCI), intracellular markers (iNOS, CD68), and the immune signals TNF, CCL2, IFN-γ, IL-6, and IL-1b. Data was analyzed using the ΔΔCT method with GAPDH used as housekeeping control and expressed as a fold change from the average ΔΔCT of the reference group (uninjected Cre-). The primer sets used are defined in Table 2.

**Table 2.**
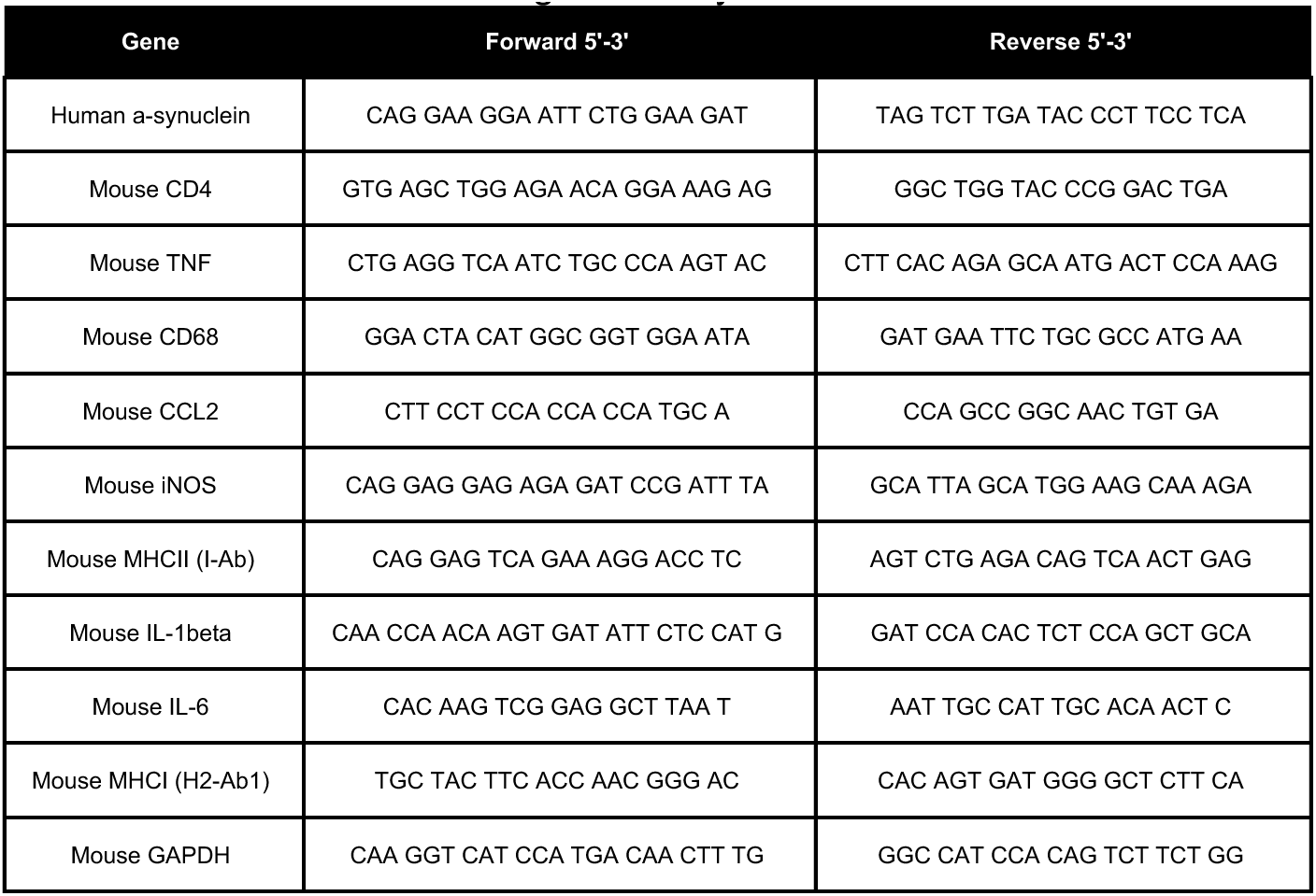
Antibodies used throughout study.

### Cohort and subject recruitment and genotyping

Individuals with PD (n=89) and healthy control subjects (n=60) were recruited through a collaboration with Dr. Nicholas McFarland at the University of Florida and samples collected as approved by the Institutional Review Boards of Emory University in Atlanta, Georgia and the University of Florida in Gainesville, Florida and in accordance with the Declaration of Helsinki. All participants were of white, European background. No participants listed Jewish religious or ethnic background. Some but not all the control subjects were spouses of PD participants. Informed consent was received in writing from all participants prior to inclusion in the study. To identify the *rs3129882* genotype of all participants, saliva was collected using Oragene DNA OG-500 kit and DNA isolated according to the manufacturers protocol (DNA genotek). Taqman SNP Genotyping Assay (Life Technologies) was used to determine genotype. Subjects provided an approximately 40mL blood sample for PBMC inducibility assay and flow cytometry. A questionnaire was administered at the time of blood collection to assess disease status, medication use, demographic background, family history, and any inflammation-related exposures. Caffeine, NSAID, and nicotine use data was collected as well.

### Human PBMC isolation, cryopreservation, and thaw

Blood was drawn via venipuncture from participants at the University of Florida into BD Vacutainer Cell Preparation Tubes (CPT) with sodium heparin (BD Biosciences Cat#362761). Following density centrifugation, the plasma layer was removed and frozen in liquid nitrogen. PBMCs were cryopreserved in 90% heat-inactivated FBS (Sigma) in 10% DMSO at a concentration of 10×10^6^ cells per 1mL. Cryovials of 1mL were frozen down in a room temperature Mr. FrostyTM (ThermoFischer Scientific) placed in a -80°C freezer for 24 hours and then shipped on dry ice to Emory University where vials were stored in liquid nitrogen until time of experiment. To thaw, 1 cryovial per participant was retrieved from liquid nitrogen and thawed at 37°C in a water bath. The thawed cell solution was then transferred into 25mL of 10% FBS in RPMI and centrifuged for 10 minutes at 300 x g at room temperature. Supernatant was removed and pellet was resuspended in pre-warmed 30 mL MACS buffer (PBS, 0.5% bovine serum albumin, 20 mM EDTA, pH 7.2) for cell counting using Trypan Blue on an Invitrogen Countess Automated Cell Counter. After counting, cells were 10 minutes at 300 x g at room temperature, supernatant was removed and pellet was resuspended in 37°C filter sterilized complete culture media for plating (RPMI 1640 media, 10% low endotoxin, 10% heat-inactivated FBS, 1mM Penicillin-Streptomycin). For human PBMC inducibility assay, 2 x 10^6^ cells were plated per condition in a 6-well culture plate. For flow cytometry, 2.5 x 10^5^ cells were plated per subject in a 96-well V-bottom plate.

### Human PBMC inducibility assay

After plating, cells were allowed to rest for 2 hours at 37°C, 5% CO2, 95% relative humidity. After resting, cells were treated with either vehicle or 100U human IFN-γ (Peprotech Cat#50813413), for 18 hours at 37°C, 5% CO2, 95% relative humidity. Cells were harvested by gently detaching adherent cells from the plate using a mini cell scraper and transferring cells into a 15mL conical falcon. Cells were centrifuged for 10 minutes at 300 x g at room temperature. Supernatant was removed and pellets were resuspended in 1mL PBS, centrifuged again for 10 minutes at 300 x g at room temperature, supernatant removed and pellets taken for RNA extraction.

### RNA extraction and cDNA synthesis and RT-PCR of Human PBMCs

RNeasy mini kit (Qiagen) was used according to manufacturer’s instructions. Briefly, 10 µL of β-mercaptoethanol was added to every 1 mL of RLT buffer. 350 µL was added to each cell pellet and cells homogenized manually with a p1000 pipette. Cell lysate was transferred to an RNAase free Eppendorf and an equal volume of 70% ethanol was added to each sample. Samples were loaded into supplier columns and centrifuged at 11,000 x g for 30 seconds and flow through discarded. 350 µL RW1 buffer was added to each column and centrifuged at 11,000 x g for 15 seconds and flow through discarded. RW1 buffer was added and columns centrifuged at 11,000 x g for 15 seconds. RPE buffer was added to columns, centrifuged for 11,000 x g for 30 seconds, repeated, and 30µL RNase-free water was added and RNA was eluted. RNA concentration was quantified, and 260/230 and 260/280 ratios recorded using a spectrophotometer.

Genomic DNA was removed from RNA samples with 1/5^th^ DNase in 25mM MgCl2 at 37°C for 30 minutes and 75°C for 10 minutes PCR cycle. Then, the reverse transcription master mix, containing 5X reaction buffer, 10mM deoxyribonucleotides, DTT, reverse transcriptase, and random primers, was added to the RNA with annealed primers. Samples were equilibrated at 25°C for 10 minutes followed by reverse transcription at 42°C for 50 minutes and reverse transcriptase inactivation at 72°C for 10 minutes. Samples were then diluted in nuclease-free water to yield an appropriate working concentration of cDNA to confidently detect gene target of interest, 10µg/µL. Gene expression was quantified using RT-PCR with SYBR Green-based chemistry. 5µL of cDNA was mixed with 35µL of RT-PCR master mix, consisting of 1.25µM forward and reverse primers^30^ obtained from Integrated DNA Technologies (Research Triangle Park), SYBR Green (ThermoFisher Scientific Cat#A46112), and nuclease-free water. This reaction mix was plated into 10µL triplicates. A QuantStudio 5 Real-Time PCR System (ThermoFisher Scientific) was used to perform the RT-PCR. Data was removed from the analysis if replicates had standard deviations >0.5. Data were analyzed using the ΔΔCT method, with data being expressed as a fold change from the average ΔΔCT of the reference group (control AA unstimulated).

### Flow Cytometry of Human PBMCs

Cryopreserved PBMCs (Control *AA* n=19, Control *GG* n=6, PD *AA* n=25, PD *GG* n=23) were thawed in 37°C water bath for ∼90 seconds and immediately transferred to 25mL of prewarmed thawing media (10% heat inactivated FBS in RPMI media). Cells were pelleted at 300 x g for 5 minutes at room temperature and then resuspended in 10mL of Macs buffer for counting. Cells were then resuspended at 1 million cells/200μL and transferred to a V-bottom plate for surface epitope labeling. Samples were centrifuged at 300 x g for 5 minutes at 4°C, washed once in PBS, re-pelleted, and resuspended in LIVE/DEAD Fixable Blue stain (ThermoFisher Scientific, #L34962) for 30 minutes at room temperature protected from the light. After live/dead incubation samples were centrifuged at 300 x g at room temperature for 5 minutes before 3 washes in 1XPBS.

Samples were then resuspended in CCR7 antibody plus FC block in BD Horizon Brilliant Stain Buffer (BD Biosciences, Cat#563794) preincubation at 37°C for 15 minutes. Samples were then spun down at 300 x g for 5 minutes at 4°C and resuspended in full antibody cocktail containing fluorophore-conjugated antibodies (Table 1) and FC block in BD Horizon Brilliant Stain Buffer (BD Biosciences, #563794). Cells were incubated with antibodies for 20 minutes at 4°C before three washes in FACS buffer containing 0.25mM EDTA, 0.01% NaN3, and 0.1% BSA in PBS. Labeled cells were fixed in 1% paraformaldehyde in a phosphate buffer for 30 minutes at 4°C. Cells were washed three times in FACS buffer and transferred to FACS tubes in FACS buffer for flow cytometry (Supplementary Figure S8). Surface markers were chosen based on the recommendations of the Human Immunology Project^55,56^.

For compensation, 1uL of antibody was added to reactive and negative AbC Total Antibody Compensation Beads (ThermoFisher Scientific, #A10497) and 0.5uL of LIVE/DEAD Fixable Blue stain was added to ArC Amine Reactive Compensation Beads (ThermoFisher Scientific, #A10346). Bead staining occurred at the same time as cell staining and was subjected to the same protocol described above without fixation (e.g., labeled beads remained in FACS buffer while cells were in fixative). Samples were run on a BD FACSymphony A3 by an experimenter blinded to the patient groups, capturing 200,000 single cells. Laser voltages were set with the assistance of SPHERO Supra Rainbow particles (Spherotech, Inc., #SRCP-35-2A) based on settings optimized during the development of this antibody panel. Compensation controls were calculated to ensure successful channel compensation prior to running samples. Raw files were analyzed using FlowJo (BD Biosciences) v.10.9.0 and gates were drawn with the assistance of fluorescence-minus-one controls.

### ATAC-seq

Tn5 transposition was performed on 1000–10,000 cells as previously described^57^. Briefly, cells were resuspended in 12.5μL 2× tagmentation DNA Buffer, 2.5μL Tn5, 2.5μL 1% Tween-20, 2.5μL 0.2% Digitonin, and 5μL H2O and incubated for 1 hour at 37°C. Next, cells were lysed with the addition of 2μL 10 mg/ml Proteinase-K, 23μL Tagmentation Clean-up buffer (326 mM NaCl, 109 mM EDTA, 0.63% SDS), and incubated at 40°C for 30 minutes. Tagmented DNA was size selected for small fragments using AMPure XP beads (Beckman Coulter, Cat#A63881) and PCR amplified with KAPA HiFi HotStart ReadyMix (Roche, Cat#KK2602) with dual indexing primers (Illumina, Cat#FC-131-2004) to generate a sequencing library. Final libraries were again size selected using AMPureXP beads, quantitated by Qubit (Life Technologies, Cat#Q33231), size distributions determined by bioanalyzer (Agilent 2100), pooled at equimolar ratios, and sequenced at the Emory Nonhuman Primate Genomics Core on a NovaSeq6000 using a PE100 run.

### ATAC-seq Data Analysis

Raw fastq reads were mapped to the hg38 version of the human genome with Bowtie2 v2.2.4^58^ and duplicate reads flagged using PICARD (http://broadinstitute.github.io/picard/) filtered based on the uniquely mappable and non-redundant reads. Enriched peaks were determined using MACS2 v2.1.0.2014061^59^ and the reads for each sample overlapping all possible peaks was calculated using the GenomicRanges v1.34.0^60^ package in R v3.5.2. Differential accessible regions/peaks were determined using DESeq2 v1.26.0^61^. All data analysis and display was performed using R v3.5.2 using scripts that are available upon request.

### Statistics

GraphPad Prism 10 software was used for all statistical analysis, with the exception of Grubbs’ test to detect outliers, which was performed using the calculator at https://www.graphpad.com/quickcalcs/grubbs1/. For mouse studies, values were compared between genotypes using unpaired 2-tailed Student’s t-test with equal standard deviation, however unpaired t-test with Welch’s correction was used when variances were not equal. To compare left and right brain hemispheres (α-synuclein virus-injected and uninjected sides, respectively) across genotypes, a two-way ANOVA was performed along with Sidak’s multiple comparison’s test. For human studies, statistical comparison among the four groups (Control *AA*, Control *AG/GG*, PD *AA*, PD *AG/GG*) was analyzed by two-way ANOVA followed by Sidak’s post-hoc multiple comparison’s test for p values. Significance for all comparisons was set at p<0.05. Data is presented throughout as mean±SEM.

## Acknowledgements

We thank the Emory Flow Cytometry Core for their assistance with the use of the LSRII, as well as the Emory Genomics Core for completing library preparations and sequencing. We thank Fredric Manfredsson for providing the AAV for stereotaxic injections and Haydeh Payami and the Tansey lab for thoughtful discussions. Partial funding from this work came from a pilot grant from Emory PD-CERC (MGT and JMB), a pilot grant from NIH/NINDS 1P50NS071669 Emory’s Udall Parkinson’s Disease Center/UL1RR025008 Atlanta Clinical and Translational Science Institute (ACTSI) (JMB and MGT), NIH RO1AI153102 (JMB), NIH/NINDS RF1NS128800 (MGT) and NIH/NINDS 5R01NS092122 (MGT and JMB). Support was also provided by Michael J Fox foundation MJFF-023296 (VJ and MGT), 1FL ADRC Alzstars P30AG066506 (VJ) and Alzheimer’s Association AARG-22-927012 (VJ). Figure 1 was created in BioRender. Jernigan, J. (2025) https://BioRender.com/r71q005.

## Author contributions

EMK and MGT conceived the study. EMK designed the study and supervised the technical staff involved in the study. EMK, JJ, CDS, JM, SLH, MKH, RLW, SDK, JC, KBM, VJ performed the various surgeries and assays. NRM provided human samples in accordance with IRB. JMB and MGT obtained funding support. JMB and CDS provided guidance and interpretation on ATAC-seq and MHCII gene expression. EMK, JJ, MGT and VJ wrote and edited the paper. EMK, JJ and VJ were responsible for all prepared figures. All authors reviewed and approved the final version of the manuscript.

## Data availability statement

The ATAC-seq datasets generated during the current study are available in the Gene Expression Omnibus repository, GSE278590. All data generated or analyzed during this study are included in this published article [and its supplementary information files].

