## Supplemental figures for "MHCII reduction is insufficient to protect mice from alpha-synuclein-induced degeneration and the Parkinson’s HLA locus exhibits epigenetic regulation"

Supplementary Figure 1.

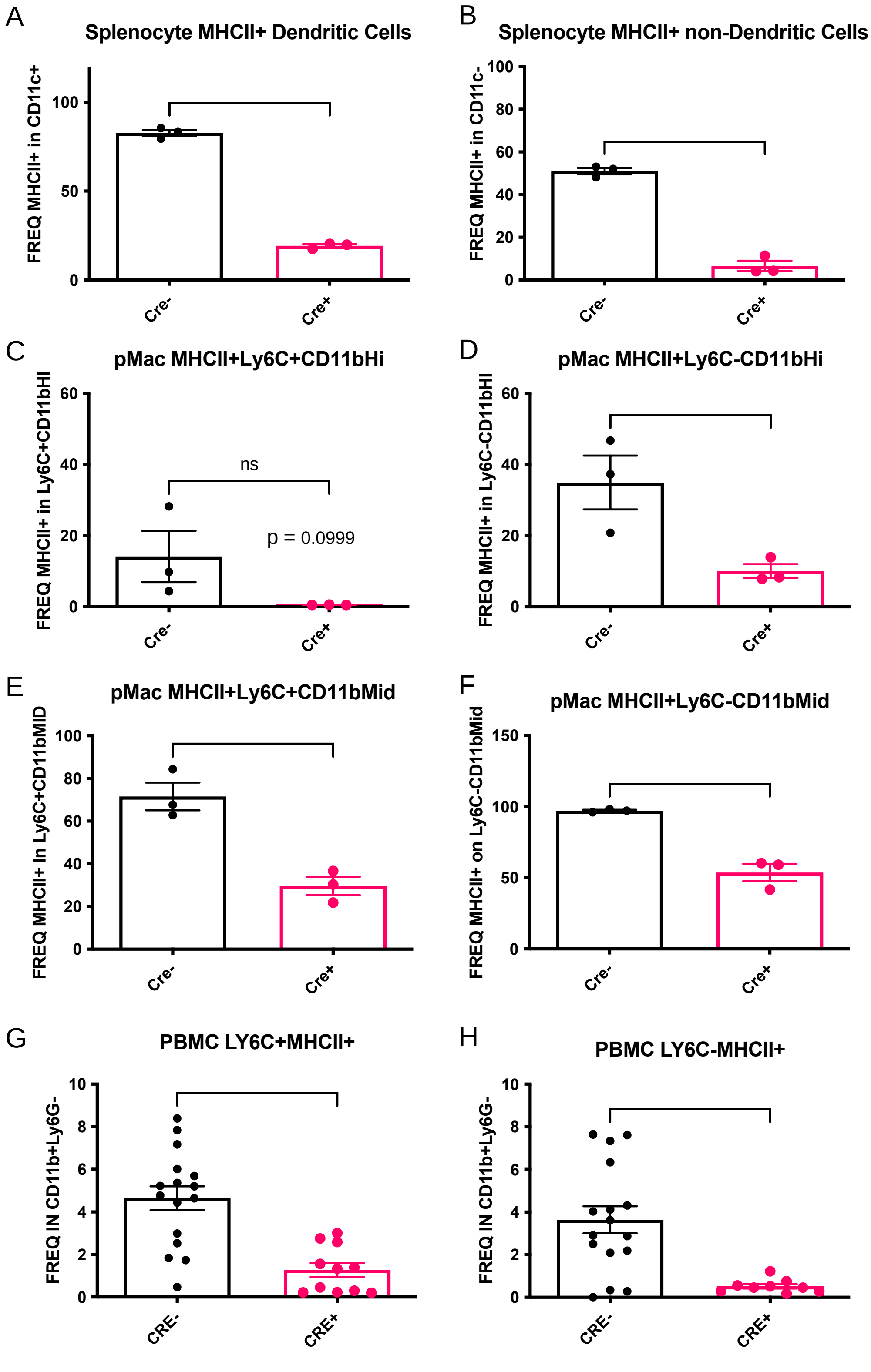


**Supplementary figure S1. Robust reduction of MHCII in myeloid cells of peripheral tissues in naïve LysMCre+I-Ab$^{fl/fl}$ mice.** Naïve mice euthanized by isoflurane-anesthetized cervical dislocation were used to verify MHCII deletion in splenocytes and peritoneal myeloid cells by flow cytometry (n=3 per genotype). The frequency of MHCII+ cells was measured by surface staining for flow cytometry on CD11b+ positively selected cells from (A,B) the spleen and (C-F) the peritoneal macrophages of naïve LysMCre-I-Ab$^{fl/fl}$ (CRE-) and LysMCre+I-Ab$^{fl/fl}$ (CRE+) mice. (G,H) MHCII was also evaluated in PBMCs of cohort 1, and monocytes, both Ly6C+ and Ly6C-, show reduced MHCII in CRE+ mice. These data indicate that CRE+ have a deletion of MHCII+ myeloid (CD11b+) cells in peripheral tissues. Data are plotted as mean +/- standard error of the mean (SEM). P-values are indicated for two-tailed t-test: ****$p<0.0001$, ***$p<0.001$, **$p<0.01$, *$p<0.05$. See supplementary figures 4-6 for gating strategy.

Supplementary Figure 2.

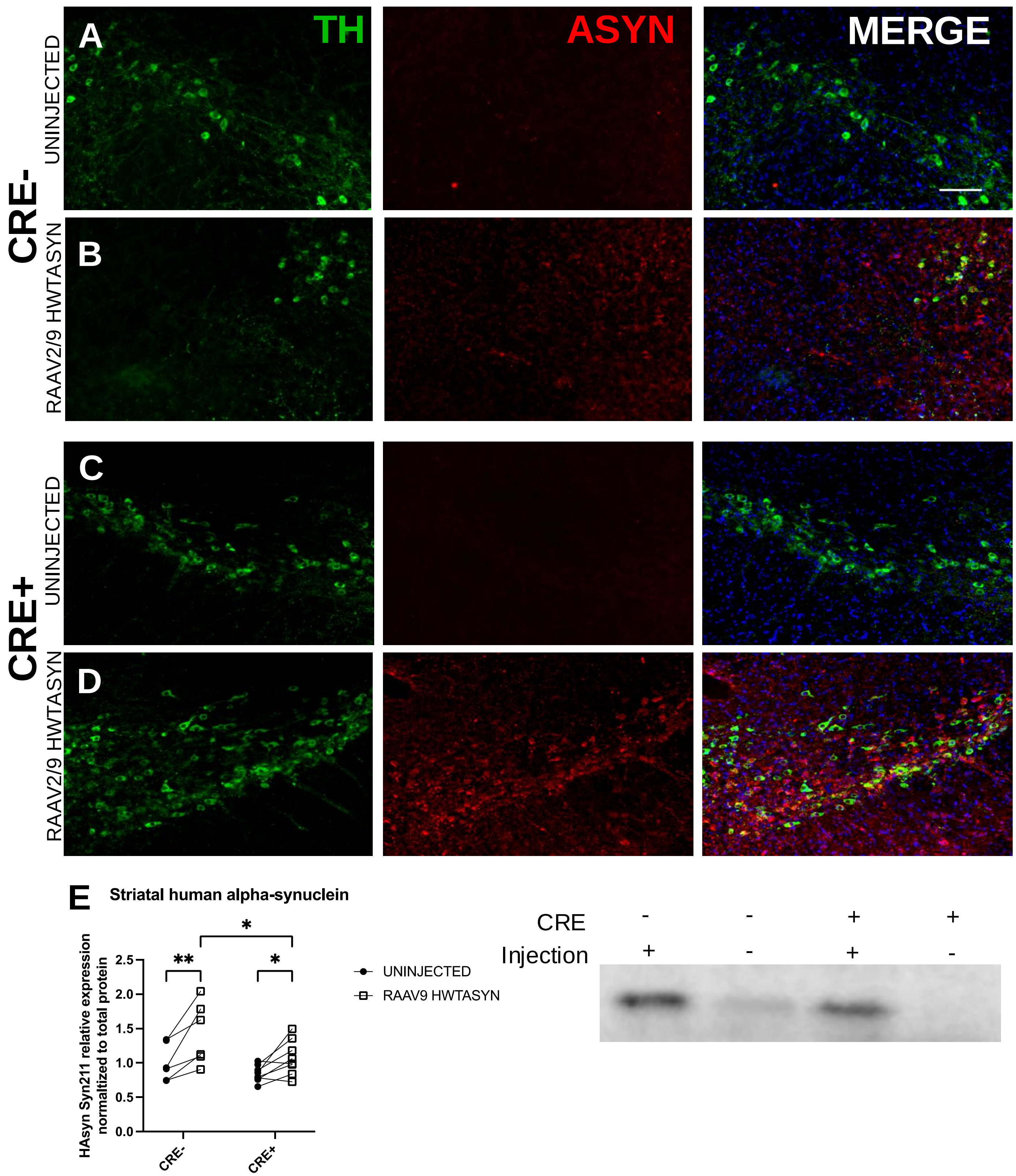


**Supplementary Figure S2. Targeting of rAAV2/9 human WT α-synuclein to the mouse substantia nigra**. (A-D) Human α-synuclein (ASYN, red) is expressed in the tyrosine hydroxylase+ (TH, green) neurons in the substantia nigra 4 months after unilateral rAAV2/9-human WT ASYN injection compared to those not injected. Images of 40µm-thick immunostained sections from LysMcre-I-Abfl/fl (CRE-; A,B) and LysMCre+I- Abfl/fl (CRE+, C,D) were taken at 20x. Scale bar = 100µm. (E) Western analysis of striatal lysates demonstrate elevated human ASYN (Syn211) in the hemisphere ipsilateral to the nigral rAAV9-human WTASYN injection compared to contralateral hemisphere. Representative blot of human α-synuclein band. **p<0.01, *p<0.05.

Supplementary Figure 3.

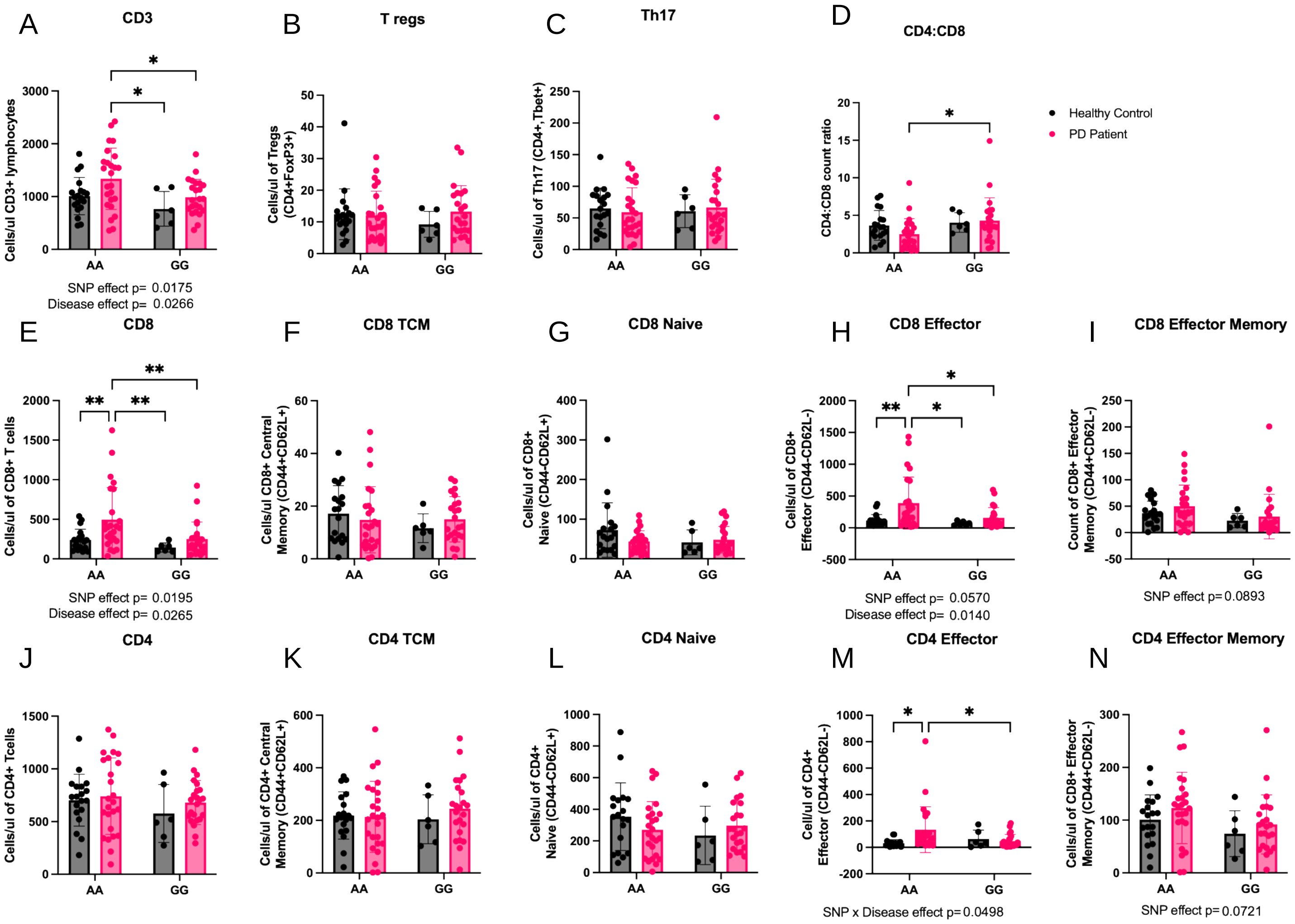


**Supplemental Figure S3. Influence of rs3129882 and PD on human T cell Immunophenotype in cryopreserved PBMCs.** Flow cytometry staining was used to determine concentrations of T cells in cryopreserved PBMC samples from healthy controls (HC) and PD patients with healthy (AA) or increased risk HLA SNP (GG). HC AA n=19, HC GG n=6, PD AA n=25, PD GG n=23. (A) Cells/µl of CD3+ T cells out of total live cells. (B) Cells/µl of CD3+CD4+CD25+CCR6+ T regulator cells out of total live cells. (C) Cells/µl of CD3+CD4+CD25+CCR6+ CCR4+ TH17 cells out of total live cells. (D) Ratio of CD4:CD8 cell count. (E) Cells/µl of CD3+CD8+ cells out of CD3+ cells. (F-I) CD8+ cell subtype cells/µl. (J) Cells/µl of CD3+CD4+ cells out of CD3+ cells. (K-L) CD4+ cell subtype cells/µl. Two-way ordinary ANOVA with Tukey correction for multiple comparisons was used to test for significant differences. Error bars represented by SEM. 2-way ordinary ANOVA with Tukey correction for multiple comparisons used to analyses the data. Trending or significant disease, SNP, and interaction effects noted below each graph. Cell counts normalized across 2 runs and calculated based on individual counting beads using the following equation: ((cell count / bead count) * (1037 beads / 10µL)*310µL). **p<0.01, *p<0.05.

Supplementary Figure 4.

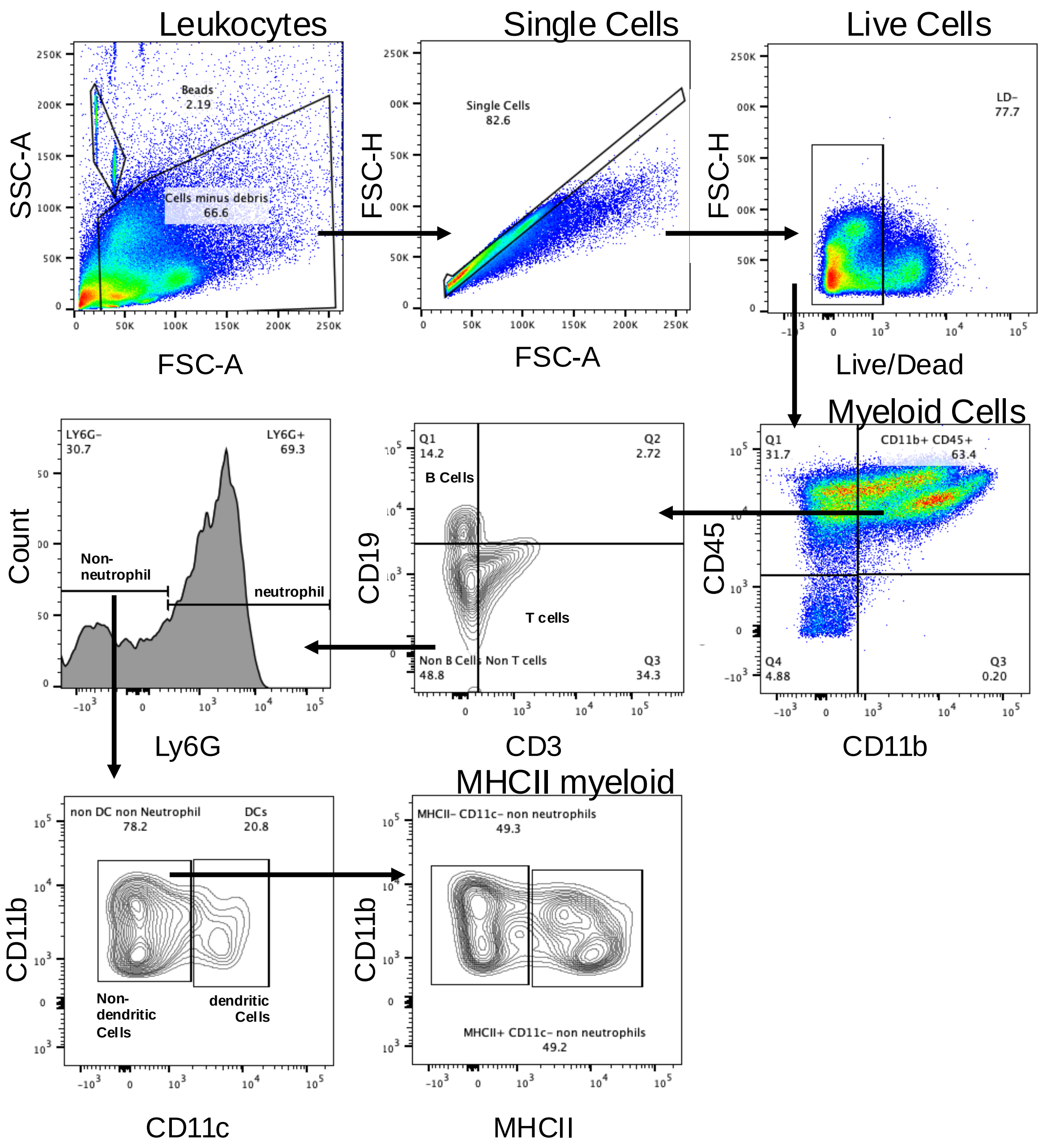


**Supplementary Figure S4. Flow cytometry gating strategies for mouse splenocytes.** Live cell population was identified as live/dead-. The myeloid immune cell population (CD11b+CD45+) was then identified and gated on CD3 (T cells) and CD19 (B cells) to further purify the myeloid population. The non-B and non-T cell cells were further gated using Ly6G+. Ly6G+ gate was used to identify neutrophils. Ly6G- cells were gated on CD11c as a marker of a subset of dendritic cells. CD11c- populations were gated on MHCII (vs CD11b) to determine the frequency of MHCII+ cells. Standardization and compensation with Sphero beads and OneComp beads was performed, and gates were drawn based on isotype antibody controls.

Supplementary Figure 5.

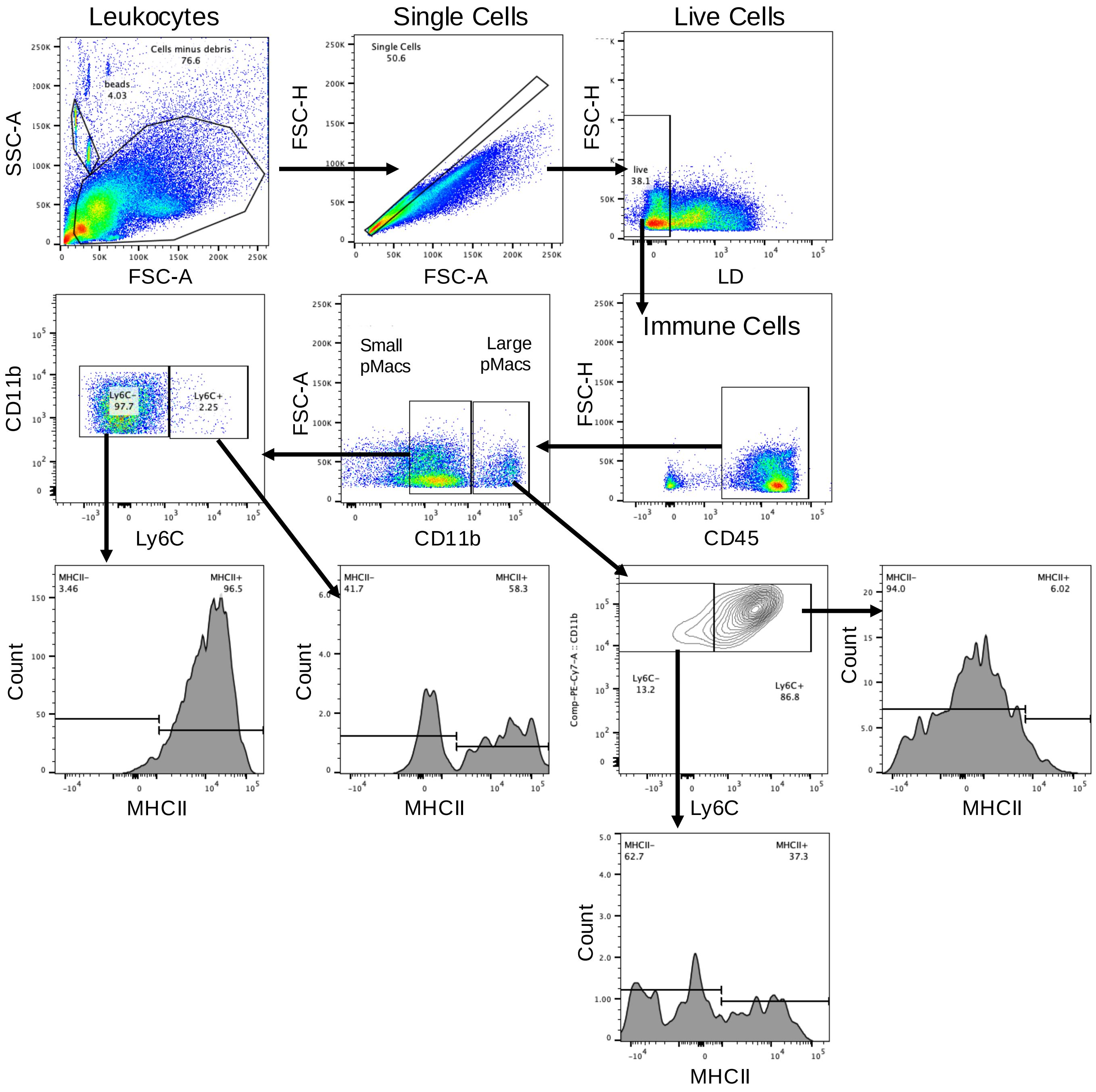


**Supplementary Figure S5. Flow cytometry gating strategies for mouse peritoneal immune cells.** Live cell population was identified as LD-. The CD45+ population (plotted versus FSC-H for clarity) was gated on CD11b (versus FSC-A for clarity), and CD11b HI (large peritoneal macrophage) and CD11b MID (small peritoneal macrophage) populations were identified. Within both the CD11bHI and MID populations, cells were gated on Ly6C to identify infiltrating (Ly6C+) and circulating (Ly6C-) cells, and both Ly6C+ and Ly6C- populations were then gated on MHCII. Standardization and compensation with Sphero beads and OneComp beads was performed, and gates were drawn based on isotype antibody controls.

Supplementary Figure 6.

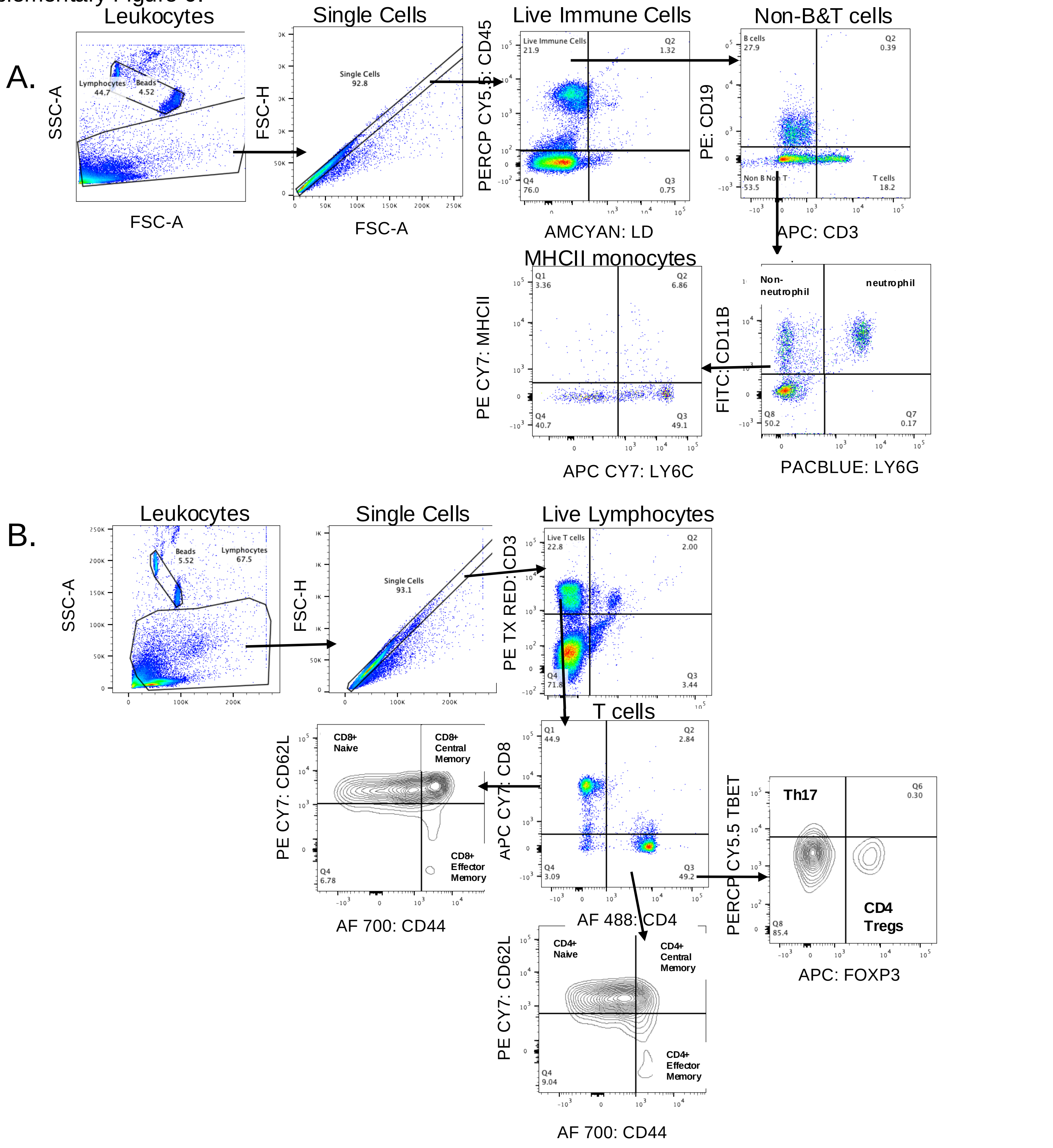


**Supplementary Figure S6. Flow cytometry gating strategies for mouse PBMC and DCLN**. (A) Live immune cells (LD-CD45+) were gated on CD3 versus CD19 to identify the T cell (CD3+CD19-), B cell (CD3-CD19+), and non-T cell, non-B cell populations (CD3-CD19-). The non-T non-B population was gated on Ly6G versus CD11b to identify neutrophils (Ly6G+CD11b+) and non-neutrophils (Ly6G-CD11b+). Non-neutrophils were then gated on Ly6C versus MHCII to identify the MHCII+ infiltrating (Ly6C+) or circulating (Ly6C-) cells. (B) The live T cell population was identified as LD-CD3+. Live T cells were then gated on CD4 and CD8. CD8+ cells were then gated on CD44 versus CD62L to identify naïve CD8 (CD44-CD62L+), central memory CD8 (CD44+CD62L+), and effector memory CD8 (CD44+CD62L-) populations. CD4+ Cells were gated on FoxP3 versus Tbet to identify Th17's (Tbet+, FoxP3-) and regulatory T cells (Tregs, Tbet-, FoxP3+). For both (A) and (B), standardization and compensation with Sphero beads and OneComp beads was performed, and gates were drawn based on isotype antibody controls.

Supplementary Figure 7.

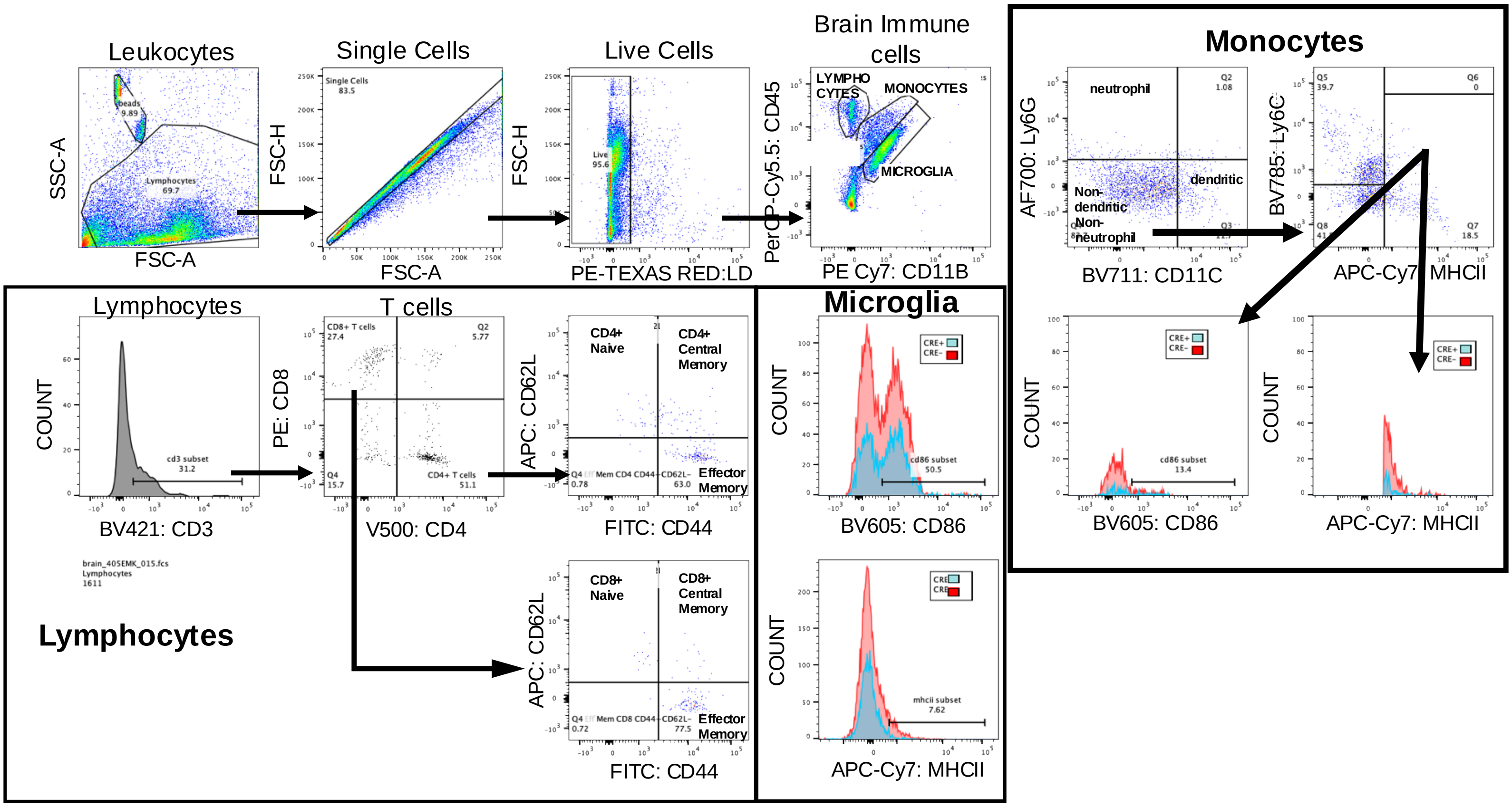


**Supplementary Figure S7. Flow cytometry gating strategy for mouse brain immune cells.** Live cell population was identified as LD-. Brain lymphocytes, monocytes, and microglia were differentiated by a CD11b versus CD45 gate. CD11b+CD45$^{low}$ cells were classified as microglia, while CD11b+ CD45$^{high}$ cells were classified as peripheral immune cells, primarily monocytes, that had trafficked to the brain. Brain lymphocytes (CD11b-CD45$^{high}$) were further gated on CD3, and the CD3+ population (T cells) was separated into CD4+ and CD8+ to characterize T cell subsets. CD44 versus CD62L gating was used to identify central memory (CD44+CD62L+), naïve (CD44-CD62L+), and effector memory (CD44+CD62L-) subsets for both CD4+ and CD8+ T cells. The monocyte population (CD11b+ CD45$^{high}$) was gated on CD11c and Ly6G to identify neutrophils (CD11c-Ly6G+) and a subset of dendritic cells (CD11c+Ly6G-). The CD11c-Ly6G- population was gated on MHCII and Ly6C to characterize monocyte activation. MHCII+Ly6C- cells were further assessed for expression of CD86. Finally, microglia (CD11b+CD45$^{low}$) were gated on CD86 and, separately, MHCII.

Supplementary figure 8.

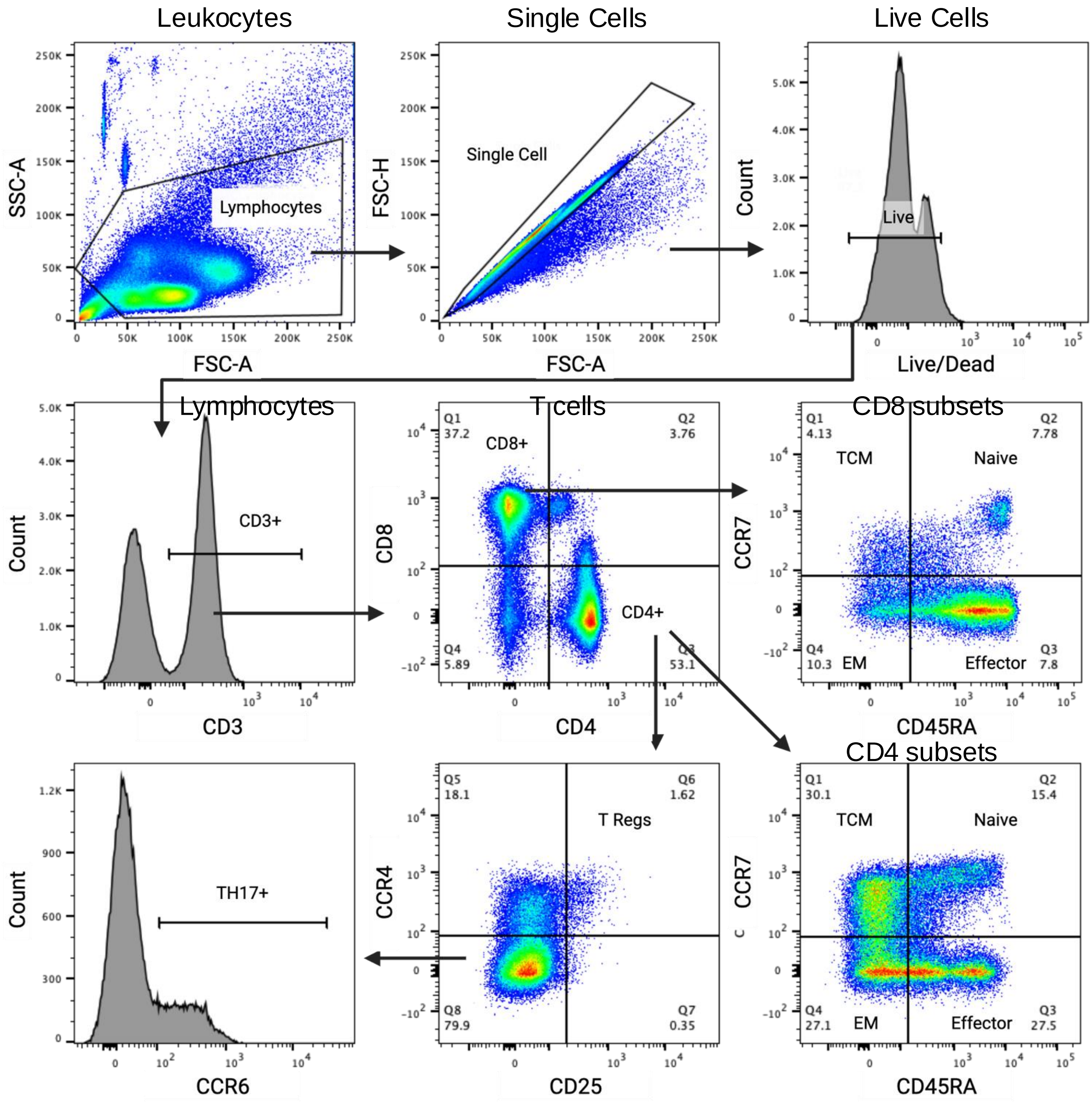


**Supplementary Figure S8. Flow cytometry gating strategy for human PBMCs.** Live single cells were identified as LD- and then gated on CD3. CD3+ cells (T cells) were gated for CD4 versus CD8. CD4+T cells were then gated on 1. CCR7 versus CD45RA and 2. CD25 versus CCR4. The following cell types were identified: Naïve CD4 T cells (CCR7+CD45RA+), central memory CD4+ T cells (CCR7+CD45RA-), effector CD4 T cells (CCR7-CD45RA+), effector memory CD4 T cells (CCR7-CD45RA-), Tregs (CD25+CCR4+), non-Tregs (CD25-CCR4-) were gated on CCR6. And Th17 (CCR6+). The same memory marker combinations were used to identify subsets of CD8+ T cells
